# Antimicrobial and Biofilm-Inhibitory Potential of Green Synthesized Silver Nanoparticles from *Lantana camara* against *Pseudomonas aeruginosa*

**DOI:** 10.1101/2025.08.19.671064

**Authors:** Achal Dharmalal Rajratna, Saurav K. Saha, Tapas K. Sengupta

**Affiliations:** Department of Biological Sciences, Indian Institute of Science Education and Research Kolkata Mohanpur, Nadia-741 246, West Bengal, India

**Keywords:** Antibacterial activity, Biofilm, Green synthesized nanoparticles, *Lantana camara*, *Pseudomonas aeruginosa*, Quorum sensing inhibition, Reactive oxygen species

## Abstract

**Background:** Emerging antimicrobial resistance poses serious threat to human well-being. Current therapeutics often fail to treat bacterial infections as pathogenic bacteria often attach to various surfaces and form biofilm. One such pathogen, *Pseudomonas aeruginosa*, can form biofilm for its survival in the presence of antibacterial agents. This study aimed to evaluate the antibacterial, antibiofilm, and mechanistic activities of *Lantana camara* leaf extract mediated silver nanoparticles (LCLE-Ag NPs) against *Pseudomonas aeruginosa*, where the extract contributes for the reduction of Ag^+^ to Ag^0^.

**Methods:** LCLE-Ag NPs were synthesized and characterized by using UV visible spectroscopy, dynamic light scattering, and electron microscopy. Antibacterial and antibiofilm activities of the synthesized NPs were assessed against *P. aeruginosa* isolates KPW.1-S1 and HRW.1-S3. Extracellular polysaccharides (EPS), and extracellular DNA (eDNA) and reactive oxygen species (ROS) generation were visualized using epifluorescence microscopy. The ability of the nanoparticles to eradicate preformed biofilm was also evaluated. The expressions of quorum-sensing genes were analyzed using semi quantitative polymerase chain reaction. Cytotoxicity was tested on human kidney epithelial-like cells.

**Results:** The synthesized LCLE-Ag NPs displayed an absorption peak at 416 nm and a size range of 30 to 35 nm. They exhibited significant antibacterial and antibiofilm activities correlating with decreased EPS and eDNA levels. LCLE-Ag NPs effectively disrupted both preformed and antibiotic-induced biofilms and such activities were found to be associated with increased ROS levels and decrease in *lasI* and *pqsA* expression. Cytotoxicity assays indicated no significant toxicity at effective concentrations.

**Conclusions:** *LCLE-*Ag NPs displayed promising antibacterial and antibiofilm activities against *P. aeruginosa*, acting through biofilm matrix disruption, ROS generation, and quorum-sensing inhibition.

## 1. Introduction

*Pseudomonas aeruginosa* (PA), Gram-negative, rod-shaped bacteria, are well-known opportunistic pathogens widely found in water, soil, plants, and animals, including humans. These pathogens are essential to study due to their inherent ability to form biofilm on different surfaces under adverse conditions as a defence mechanism and for its survival, making it clinically relevant. *Pseudomonas aeruginosa* causes secondary infections in immunosuppressed individuals as well as patients who have cystic fibrosis (1,2), chronic wounds (3), mastitis (4), endometritis (5), and thermal wounds (6). As stated in reports by the Centers for Disease Control (CDC) and Prevention experts, it is estimated that about 65% of human bacterial infections are linked to biofilms, highlighting their impact on health and the challenges they pose for treatment (7,8).

*Pseudomonas aeruginosa* isolates are generally used as model organisms for biofilm-related studies as they are avid biofilm formers, causing chronic infections (8). Besides the site of infections, PA can grow as biofilm on a variety of surfaces, including medical implants and catheters, making it clinically more vital. Biofilm also helps bacteria avoid phagocytosis and defend against different constituents of the host’s adaptive and innate immune systems, making it resistant to antimicrobial treatments (8). It is reported that antimicrobial resistance (AMR) was heightened in biofilm cells compared to planktonic cells (9,10). Thus, the increasing mortality and antimicrobial resistance of *Pseudomonas aeruginosa* highlight the need to develop novel therapeutics against the bacteria for combating these infections.

Different novel strategies are being developed against *Pseudomonas* infections in recent times due to their ability to develop new resistance mechanisms to existing strategies. Newer strategies such as anti-quorum sensing molecules, synthetic small organic molecules (11), antibiofilm peptides (12), matrix-degrading enzymes (13), phage therapy (14), and CRISPR-Cas9 (15), photodynamic therapies (16), vaccines (17), nanoparticles and biomaterials (18) are in use to enhance treatment effectiveness against infections associated with biofilms. Nanoparticles, in particular, have been an emerging tool in the recent field of therapeutics.

Nanotechnology-based tools have several advantages over different novel therapeutics, such as a high surface area to volume ratio, reactivity, and stability (19). These nanotechnologies have shown a positive impact in the field of biomedicine and pharmaceuticals, due to their effectiveness in reducing infections with conventional drug administration methods (20). Synthesis of nanoparticles through chemical synthesis involves an energy-intensive process with harsh chemical involvement, which can be overridden by green synthesis of nanoparticles, as this method offers non-toxic plant extract involvement, which acts as a reducing agent (21). The green synthesized nanoparticles can be produced on a large scale with an eco-friendly approach. The involvement of biological components in a reducing and capping agent avoids high energy and pressure needs and, simultaneously less hazardous than chemically synthesized counterparts (22,23). Among different types of nanoparticles, green synthesized metal-based nanoparticles have exhibited good potential as antimicrobial agents (24), antifungal agents (25), and also as anticancer agents (25,26).

Silver nanoparticles have been recognized for their potent antibacterial properties, effectively targeting both Gram-negative and Gram-positive bacterial strains (27,28). Silver nanotechnology can potentially inhibit bacterial infections by following the mode of action as cell membrane damage, producing different types of reactive oxygen species (ROS), and inhibiting enzymes (29,30). Such green synthesized silver nanoparticles are known to be effective, and they have many potential applications (31).

*Lantana camara*, an invasive, adaptable weed native to the American tropics, is a flowering plant of the Verbenaceae family. It has spread across the world, including India. *L. camara* has been used in traditional medicine to treat various diseases (32). Different studies have found that *L. camara* leaves display bactericidal, insecticidal, fungicidal, nematocidal, antioxidant, and anticancer properties (33,34). Traditionally, various parts of this plant, like flowers, leaves, and roots, have been utilized in herbal medicine (35). Bioactive compounds present in the plant have been found to have antibacterial activity. As only a tiny fraction of plant species has been thoroughly investigated for their therapeutic benefits, deep research into the potential of *L. camara* as an antimicrobial agent may lead to the discovery of novel treatments that are both effective and less toxic than current chemotherapeutic agents.

Metallic nanoparticles have been investigated over the past few years, and it has demonstrated the potential against clinically relevant pathogens, including *P. aeruginosa* (36,37). Yadav et al. showed that silver nanoparticles are quite effective in inhibiting *PA* biofilms (38). Though metal-based nanoparticles have promise, their toxicity limits their uses (39). Thus, green synthesis approaches have shown a promising alternative for PA infection due to their eco-friendly, less toxic and biocompatible nature (21). Various biosynthesized nanoparticles were also found to be effective against biofilm formation, as well as the virulence genes of PA (40). *Lantana camara* conjugated silver nanoparticles were also found to have an antibacterial effect on various bacteria (41). Although these studies confirm the antibacterial potential of the *Lantana camara* conjugated silver nanoparticles, their antibiofilm potential has not yet been explored in depth.

Hence, in the current study, *Lantana camara* leaf extract conjugated silver nanoparticles (LCLE-Ag NPs) were green synthesized and characterized. The synthesized nanoparticles were evaluated for their antimicrobial and antibiofilm properties against two strains of *Pseudomonas aeruginosa,* KPW.1-S1 and HRW.1-S3, which were isolated from contaminated water sources and found to express various virulent genes (42). As biofilm formation depends on the extracellular matrix components such as exopolysaccharides and extracellular DNA, the alterations of their abundance due to nanoparticle were examined. Nanoparticle-mediated antibacterial activities are often associated with oxidative imbalance. Therefore, reactive oxygen species (ROS) generation was analyzed to find any probable contribution of oxidative stress associated with the antibacterial activity of LCLE-Ag NPs. Moreover, because sub-MIC concentrations of antibiotics, including gentamicin (42), are known to induce biofilm formation, the efficacy of the nanoparticles to reduce or mitigate preformed biofilms induced by gentamicin was evaluated.

Quorum sensing plays a pivotal role in biofilm formation of *Pseudomonas aeruginosa*. The *lasI* (gene name for acyl-homoserine-lactone synthase) and *pqsA* (gene name for anthranilyl-CoA synthetase) are two important regulator genes for *P. aeruginosa* quorum sensing, which result in the enhanced expression of virulence factors and biofilm formation. In the Las quorum sensing system, *lasI* synthesizes 3-oxoC12-HSL which in turn activates other biofilm related genes. Consequently, *pqsA* is part of the PQS (*Pseudomonas* Quinolone Signal) system, working alongside the Las system during biofilm formation. It regulates biofilm physiology and development and is responsible for eDNA production (43,44). Given the central role of these two genes in the regulating biofilm, the expression of *lasI* and *pqsA* in the presence of nanoparticles was analyzed to gain molecular insights into the working mechanism of LCLE-Ag NPs.

## 2. Materials and Methods

### 2.1. Bacterial strains of *Pseudomonas aeruginosa*

The *Pseudomonas aeruginosa* strains KPW.1-S1 and HRW.1-S3 were isolated from Kolkata Port water and Haldi River water, respectively (45) and further identified by 16S rRNA sequencing (NCBI Accession numbers FJ897721 and FJ897723, respectively). The strains were deposited at the Microbial Type Culture Collection and Gene Bank (MTCC), Chandigarh, India, under the catalogue numbers MTCC 10087 and MTCC 10088 (45).

For growth, *P. aeruginosa* KPW.1-S1 and HRW.1-S3 were inoculated and grown in a modified Bushnell-Haas (BH) medium supplemented with 2% glucose as the carbon source (46).

### 2.2 Cell Culture of HEK293T cells

The HEK293T (human kidney epithelial-like) cell line was obtained from the National Centre for Cell Science, Pune, India. The cell line was maintained in Dulbecco’s Modified Eagle Medium (DMEM) High Glucose (HiMedia, India), supplemented with 10% Fetal Bovine Serum (FBS) (HiMedia, India), 10000 U/mL Penicillin, and 10 mg/mL Streptomycin antibiotic solution (HiMedia, India). Cells were maintained at 37°C in a humidified environment with 5% CO₂ and 95% air to ensure optimal growth conditions (35).

### 2.3. *Lantana camara* leaf extract (LCLE) preparation

The leaf extract was prepared following the protocol described by Pal et al. (35), with slight modifications. Healthy *L. camara* leaves were collected from the Indian Institute of Science Education and Research, Kolkata campus (22°57′50″N, 88°31′28″E). The leaves were taxonomically verified by the Central National Herbarium, Botanical Survey of India (West Bengal, India), with a voucher reference number IISER/AP/01. Then they were thoroughly washed with water and then shade-dried for 48 hours. The dried leaves were powdered using a mortar and pestle for subsequent processing.

Leaf powder was extracted with absolute ethanol (Merck) at a 1:10 w/v ratio at 37 °C for 18 hours in a shaker. The extract was filtered, concentrated by a rotary evaporator, reconstituted in molecular-grade ethanol (Merck, Germany), sterilized through a syringe filter (HiMedia) of pore size 0.22-μm, and stored at 4 °C.

### 2.4. Synthesis of silver nanoparticles

A 2 mg/mL solution of ethanol-based leaf extract of *L. camara* dispersed in water and a 3 mM silver nitrate solution were mixed in a conical flask in a 2:1 ratio. The silver nitrate solution was added drop wise over the course of one hour. It was then incubated in a hot water bath at 60 °C for three hours until the solution turned brownish-yellow. The resulting mixture was centrifuged (13,000 rpm, 10 minutes), and the pellet was resuspended in water and stored at 4 °C for further use (41).

Based on the 2:1 ratio, the maximal concentration of plant extract of *L. camara* in synthesized nanoparticles would be 1.33 mg/mL, assuming complete utilization of the extract while nanoparticle formation.

### 2.5. Characterization of Nanoparticles

#### 2.5.1. Ultraviolet-visible spectroscopy of LCLE-Ag NPs

The synthesized nanoparticles were analyzed using a UV-Vis spectrophotometer (Beckman Coulter DU 730). The analysis was conducted in the wavelength range of 200–800 nm, utilizing continuous scanning mode to capture detailed spectral data. Distilled water was employed as a blank.

#### 2.5.2. Dynamic Light Scattering and Zeta Potential of LCLE-Ag NPs

The polydispersity index (PDI), average particle size, and zeta potential were measured using a Zetasizer (Nano ZS, Malvern) using DLS. The samples were prepared by dispersing them with sonication and diluting them with MiliQ water as required.

#### 2.5.3 Scanning Electron Microscopy and Energy-Dispersive X-ray spectroscopy of LCLE-Ag NPs

The morphological characteristics and elemental composition of the synthesized LCLE-Ag NPs were studied using field emission scanning electron microscopy (FESEM) integrated with energy-dispersive X-ray spectroscopy (EDX) (Supra 55 VP, Carl Zeiss). The nanoparticle was drop-cast on cover slips, then mounted on aluminium stubs for SEM analysis, and *L. camara* powdered leaf extract was directly mounted on stubs for EDX analysis. The samples were subjected to sputter coating and subsequently analyzed under the electron microscope.

#### 2.5.4. Transmission Electron Microscopy of LCLE-Ag NPs

Transmission electron microscopy (TEM) (JEM-2100 Plus, JEOL) was performed to determine the size of the nanoparticles. Samples were drop-cast on a carbon-coated copper grid, which was then left to air-dry overnight. The samples were desiccated and visualized under TEM at an accelerating voltage of 300 kV.

#### 2.5.5. Atomic Force Microscopy of LCLE-Ag NPs

To prepare the LCLE-Ag NP sample for examination using Atomic Force Microscopy (AFM) (Cypher asylum Research AFM, Oxford Instruments), the nanoparticle was drop-cast on a silicon wafer and was placed on a tissue paper-lined petri dish, allowed to air dry in a laminar flow hood. Afterwards, the air-dried silicon wafer was desiccated and then examined under the microscope.

#### 2.5.6. Attenuated Total Reflectance (ATR) FTIR of LCLE-Ag NPs

ATR was used to examine the presence of functional groups in the nanoparticles and *L. camara* leaf extract powdered samples. ATR was performed using a PerkinElmer Spectrum RX1 spectrophotometer in the scanning range of 500-4500 cm^−1^.

### 2.6. Minimum Inhibitory Concentration (MIC) assay

The MIC assay was used to evaluate the minimal inhibitory concentration of LCLE-Ag NPs against two different strains of *Pseudomonas aeruginosa*, namely KPW.1-S1 and HRW.1-S3. Various concentrations of LCLE-Ag NPs (5- 10- 20- 50- 100 µg/mL) along with the maximal concentration of 1.33 mg/mL of LCLE were prepared and added to bacterial cultures. The individual bacterial cultures treated with nanoparticles and LCLE were incubated in a shaker (37°C, 150 rpm) along with their respective blanks. Following 18 hours of incubation, the optical density (OD) was measured using a UV-Vis spectrophotometer (Schimadzu) at 600 nm.

### 2.7. Quantification of *Pseudomonas aeruginosa* Biofilm

The biofilm load assay was performed on 24-well plates coated with polystyrene (NUNC). 3 × 10^6^ cells/mL of *Pseudomonas aeruginosa* KPW.1-S1 and HRW.1-S3 were individually cultured along with the different concentrations of LCLE-Ag NPs and also in the presence of 1.33 mg/ml LCLE in an orbital shaker (37 °C, 150 rpm). After 24 hours of incubation, the optical density of the planktonic cells was measured at 600nm using a UV-Vis spectrophotometer. The wells were rinsed twice with Phosphate Buffered Saline (PBS) to remove any excess planktonic cells, followed by staining the wells with 0.1% Crystal violet (CV) for 45 minutes. Post incubation, the CV was discarded and the wells were washed twice with PBS again. Then, 1 mL of 30% acetic acid was added to each well, and the plates were kept in the shaker (37 °C, 150 rpm) for 30 minutes to solubilize the stain. The optical density of the solubilized crystal violet was measured using a plate reader at 600 nm (BioTek ELx800).

### 2.8. Confocal Laser Scanning Microscopy for Biofilm Visualization

*Pseudomonas aeruginosa* KPW.1-S1 and HRW.1-S3 biofilm formation was further studied using confocal laser scanning microscopy (CLSM) to evaluate the impact of maximal concentration of LCLE and different doses of LCLE-Ag NPs. Biofilms were grown on coverslips in 24-well plates with respective treatment doses and incubated at 37 °C (150 rpm). Following 24 hours of incubation, the coverslips were carefully washed twice with PBS and stained with 0.005% (w/v) acridine orange for 5 minutes. Following staining, the coverslips were rinsed twice with PBS and mounted onto clean glass slides using 5% glycerol. Transparent nail polish was used to seal the edges of the coverslip. The samples were examined using a Nikon confocal microscope with an excitation wavelength of 488 nm, and emission was detected in the 293–586 nm range (42).

### 2.9. Scanning Electron Microscopy of *Pseudomonas aeruginosa* biofilm

The *Pseudomonas aeruginosa* KPW.1-S1 and HRW.1-S3 biofilm was grown on a glass coverslip in 24-well plates. After the 24-hour incubation period, each coverslip was washed twice with PBS to remove any planktonic cells. The coverslips were then immersed in 2.5% glutaraldehyde for 2 hours at 4 °C in the dark to fix the biofilm structure. Then they were washed thrice with PBS. Next, the coverslips were treated with 0.1% osmium tetroxide for 45 minutes at 4 °C in the dark. After another series of three PBS washes, the coverslips were subjected to a dehydration process using an alcohol gradient from 30% to 100%, with each concentration applied for 10 minutes. Finally, the coverslips were dried under a vacuum in a desiccator and examined in Scanning Electron Microscopy using Carl Zeiss smart software for detailed analysis of the biofilm structure (42).

### 2.10. Visualization and estimation of Extracellular polysaccharide and DNA (eDNA)

The two important extracellular matrix components, exopolysaccharides and extracellular DNA of *Pseudomonas aeruginosa* KPW.1-S1 and HRW.1-S3 biofilms grown absence and presence of different doses of nanoparticles were visualized and analyzed. The exo-polysaccharide and eDNA were visualized by dual staining of Concanavalin A (Con A) (25 μg/mL in PBS) and Propidium Iodide (PI) (0.2 μg/mL in PBS) using Olympus-IX83 fluorescence microscope. The coverslips with biofilm were washed twice with PBS and stained with Con A for 15 minutes, followed by staining with PI for 10 minutes. The coverslips were then washed with PBS and mounted on a glass slide with 5% glycerol and imaged under the fluorescence microscope. The Con A and PI were excited at 495 nm and 493 nm with emission peaks at 519 nm and 636 nm, respectively. The fluorescence intensities of the images were measured using ImageJ software and graphs were plotted accordingly.

### 2.11. Biofilm eradication assay

Similarly, a pre-formed biofilm load assay was performed on *Pseudomonas aeruginosa* KPW.1-S1 and HRW.1-S3, to estimate the biofilm eradication potential of the LCLE-Ag NPs on the formed biofilm. The biofilm was first grown for 24 hours. After this initial growth period of 24 hours, the media with planktonic cells were carefully removed and replaced with fresh BH 2% glucose media with different concentrations of LCLE-Ag NPs. The rest of the steps were followed according to the biofilm load assay protocol (Section 2.7).

### 2.12. Morphological study of the effect of LCLE-Ag NPs on gentamicin induced preformed biofilm

To check the effect of the nanoparticle on this gentamicin induced biofilm, CLSM was performed. Biofilm was grown on coverslips in the presence of 1.5 μg/mL of gentamicin in BH 2% glucose media in a shaker for 24 hours. Post-incubation, the media was replaced with fresh media containing different doses of nanoparticles. Post incubation the coverslips were washed twice with PBS, followed by a 5-minute staining in 0.005% acridine orange (w/v) solution in the dark. The stained coverslips were washed with PBS and then mounted on a glass slide with 5% glycerol, and sealed with nail polish. The samples were visualized using a Nikon confocal microscope with an excitation wavelength of 488 nm, and emission was detected in the 293–586 nm range.

### 2.13. Reactive Oxygen Species (ROS) production assay

ROS generation was measured for both the strains using an oxidation-sensitive fluorescent probe, dichlorodihydrofluorescein diacetate (DCFH-DA) (40). *P. aeruginosa* KPW.1-S1 and HRW.1-S3 Biofilms were grown on coverslips in a 24-well plate and incubated at 37 °C (150 rpm) for 24 hours. After incubation, the coverslips were gently washed twice with PBS and stained with 10μM DCFH-DA for 30 min. The prepared coverslips were mounted on a glass slide using 5% glycerol and sealed with transparent nail polish. The samples were visualized using an Olympus -IX81 fluorescence microscope, with a blue excitation filter (460-495nm) and emission filter (510-550nm). The intensity was analyzed using ImageJ software.

### 2.14. Determination of mRNA level expression of *lasI* and *pqsA* in PA KPW.1-S1 biofilm cells

Semi quantitative polymerase chain reactions were performed to check the expression of quorum-sensing genes, *lasI* and *pqsA* in absence and presence of different doses of nanoparticles. The bacterial biofilm was grown in different conditions and harvested for RNA isolation. The RNA was isolated following the protocol described in Pal et al. (35) with minor modifications. The RNA was quantified and further used for the cDNA synthesis using PrimeScript cDNA synthesis kit (TaKara, Japan) according to the manufacturer’s protocol. The synthesized cDNA was then used as a template for carrying out PCR of the genes (Table 1). The PCR reaction was carried 25 cycles for 16s, 38 cycles for both *lasI* and *pqsA*. The amplified PCR products were run on agarose gel electrophoresis and band quantification was done using GeneTools software. The intensity of the bands were normalized using their respective housekeeping bands of the same experimental condition and fold change was plotted.

**Table 1:**
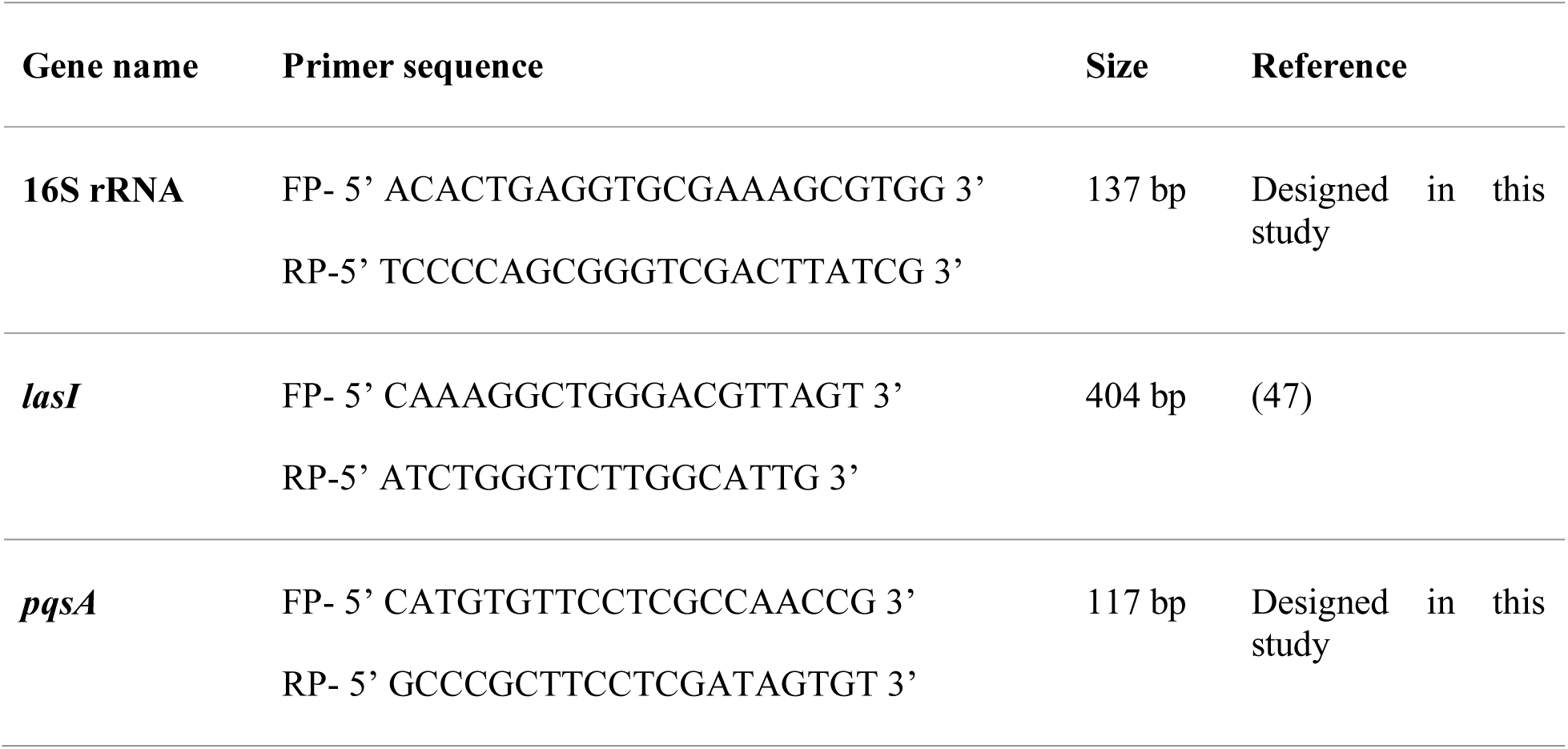
List of primers used in this study.

### 2.11. Cell Viability Assay

The cytotoxicity of LCLE-Ag NPs, along with their control (equivalent Ag concentration), was tested on HEK293T cells using a standardized 3-(4,5-dimethylthiazol-2-yl)-2,5-diphenyltetrazolium bromide (MTT) colorimetric assay protocol (35). Briefly, human epithelial kidney like (HEK293T) cells were seeded in a 96-well culture plate with 5000 cells/well and allowed to adhere overnight under standard conditions (37 °C, 5% CO_2_). Cells were then treated with 10- 20- 50- 100 μg/mL LCLE-Ag NPs and their respective silver concentrations, along with an untreated control, for 24 hours. Following 24-hour treatment, the MTT assay was performed by replacing spent media with fresh medium containing 0.5 mg/mL of MTT solution (SRL, India). After 4-hour incubation in darkness at (37 °C, 5% CO_2_), the formazan conversion reaction was terminated by carefully removing the MTT solution and dissolving the crystalline products with 100 μL dimethyl sulfoxide per well and then incubated for 30 minutes in the dark at 37 °C. Then, the absorbance was taken in a microplate reader at 595 nm. The cytotoxic effect of the LCLE-Ag NPs and AgNO_3_ was expressed as cell viability percentage compared to the untreated control.

### 2.12. Statistical analysis

The statistical analysis was carried out using GraphPad Prism 9.5.1, with results expressed as mean ± standard error. A two-tailed Student’s t-test was employed to evaluate differences between group means, with statistically significant differences denoted by asterisks (*P<0.05; **P<0.01; ***P<0.001).

## 3. Results

### 3.1. Characterization of green synthesized silver nanoparticles

#### 3.1.1. UV-Visible spectral analysis

UV-Visible spectroscopy was performed to check the absorbance band of the prepared LCLE-Ag NP. The distinct peak between 300-500 nm of the prepared nanoparticles revealed the Surface Plasmon Resonance (SPR) peak, indicating the formation of the nanoparticle, with the maximum absorbance at 416 nm (Figure 1).

**Fig. 1.**
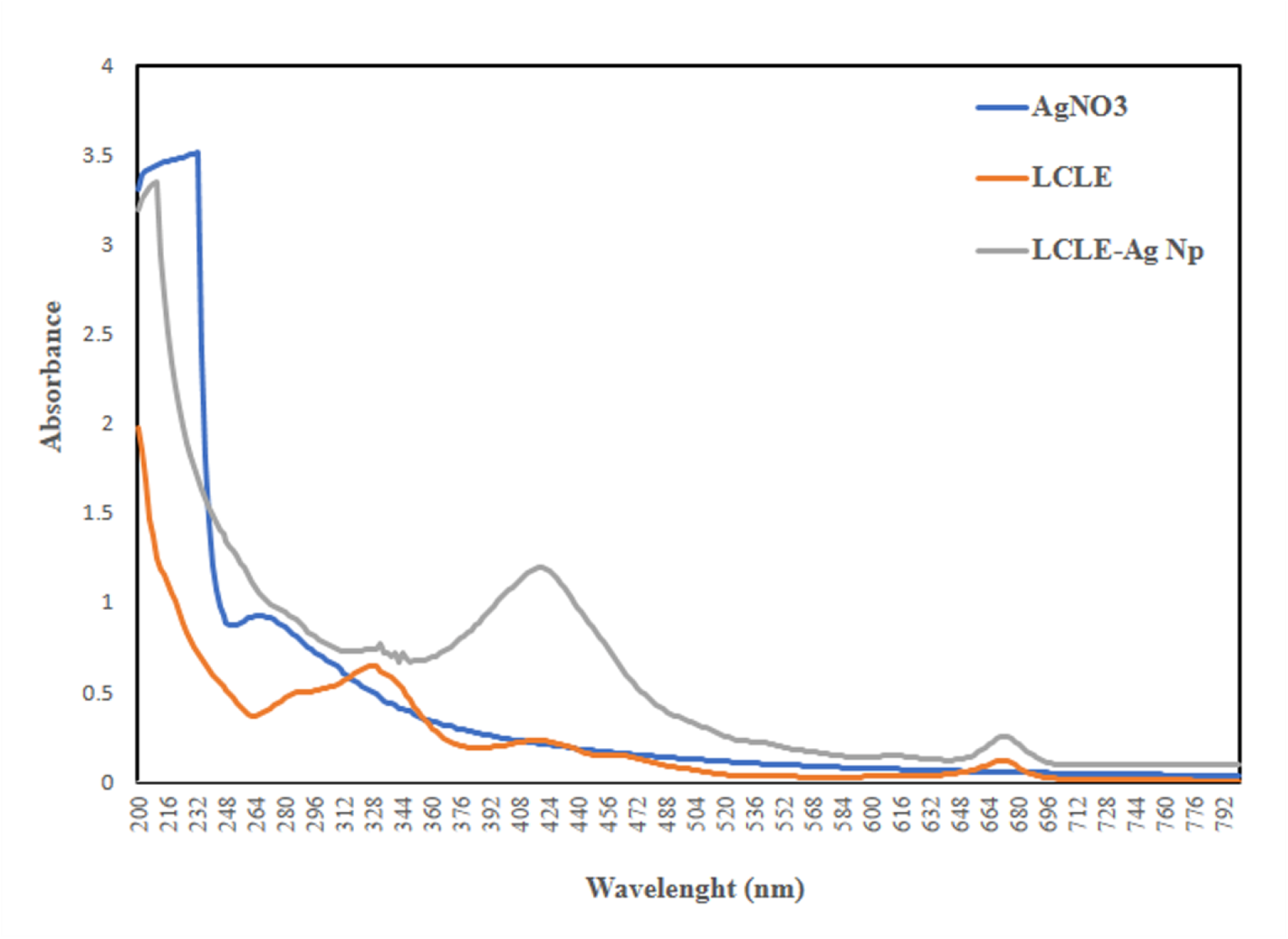
UV–vis absorbance spectra of AgNO₃ solution (blue line) and *L. camara* leaf extract (Orange line), and LCLE-Ag NPs (grey line)

#### 3.1.2. Dynamic Light Scattering analysis

Dynamic Light Scattering was studied to examine the average size and PDI of LCLE-Ag NPs. The average size of the nanoparticles determined by the size analyzer was 35 nm (Figure 2a). The synthesized nanoparticles showed a PDI of 0.3, signifying a homogenous population of nanoparticles. Overall, the result suggests that the synthesized nanoparticle is homogeneous, with a size of around 20-40 nm.

**Fig 2.**
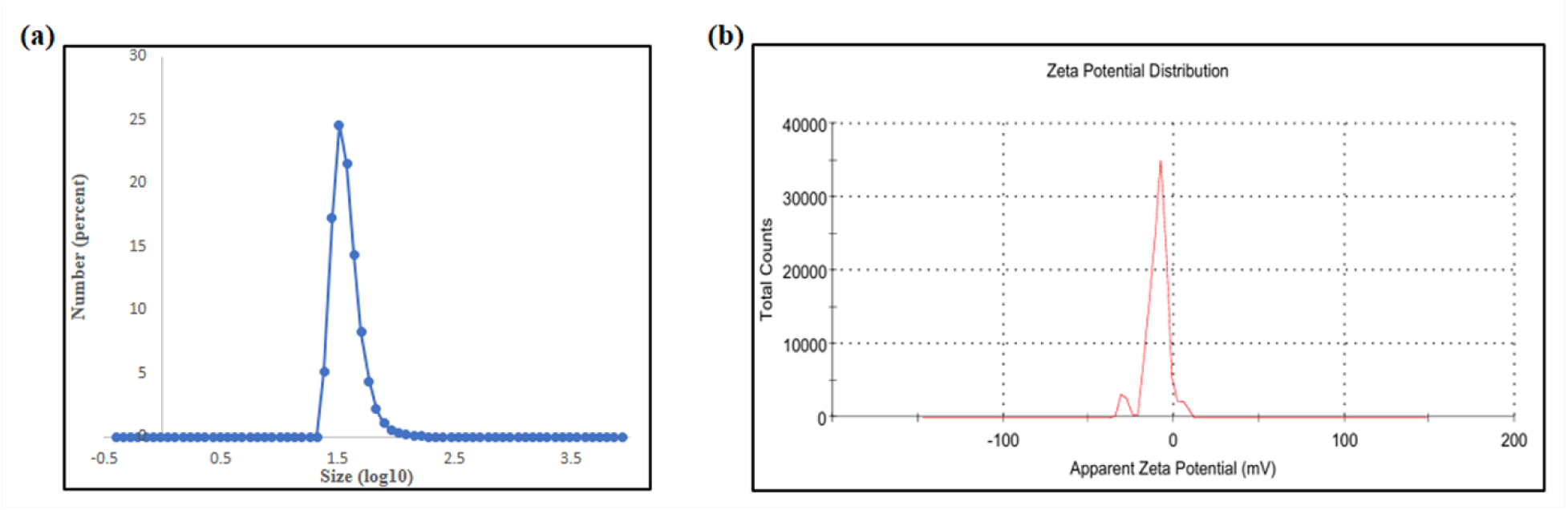
Particle size and zeta potential analysis of green synthesized LCLE-Ag NPs (a) Particle size analysis, (b) Zeta potential analysis of LCLE-Ag NPs

#### 3.1.3. Zeta potential analysis

The surface charge of the synthesized LCLE-Ag NPs was determined using a zeta sizer. The graph displays the corresponding average zeta potential of -9.53 mV (Figure 2b). This zeta potential value implies the stability of the synthesized nanoparticles, as the negatively charged nanoparticles will exhibit a repulsive force, resulting in enhanced stability.

#### 3.1.4. Atomic Force Microscopy analysis

AFM observations corroborate the obtained morphological results of synthesized nanoparticles. The AFM images revealed that LCLE-Ag NPs exhibited a nearly spherical shape with some degree of agglomeration from lateral and horizontal views (Figure 3a,b). The height and latitude studied in the analysis support the data, suggesting the spherical shape of the prepared nanoparticle.

**Fig 3.**
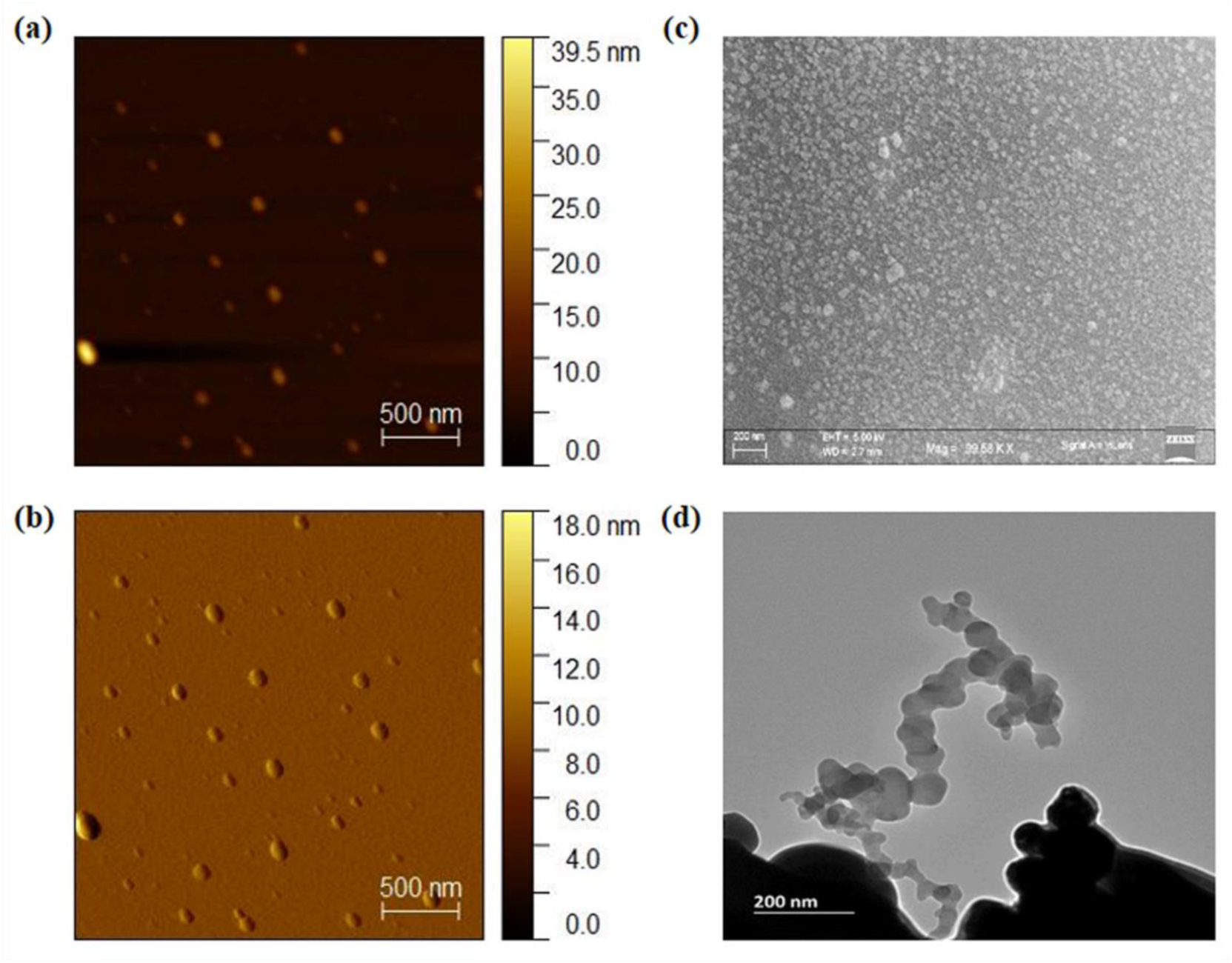
Morphological and characteristic analysis of green synthesized LCLE-Ag NPs (a) AFM height, (b) AFM amplitude, (c) SEM, (d) TEM

#### 3.1.5. Scanning Electron Microscopy and Energy Dispersive X-ray Spectroscopy analysis

Scanning Electron Microscopy was performed to depict the morphology of the synthesized nanoparticles (Figure 3c). The morphology of the prepared nanoparticles was observed to be nearly spherical with fair agglomeration. The topographical analysis revealed that the average particle size is 26 nm.

Energy Dispersive X-ray Spectroscopy (EDX) analysis was used to analyze the elemental composition of the synthesized LCLE-Ag NPs (Supplementary Figure S1a). The graph indicates a significant peak for Silver with a weight percentage of nearly 70%. The EDX analysis also confirmed the formation of LCLE-Ag NPs.

The EDX of *L. camara* leaf extract was done to analyze elemental composition (Supplementary Figure S1 b). Quantitative analysis elucidates that Cl, O, Ca, K, Si, and Ag elements are present in the leaf extract. Out of these, O and K carried out the majority weight percentages of 78.53% and 11.26%, respectively, with traces of other elements.

#### 3.1.5. Transmission Electron Microscopy

The Transmission Electron Microscopy images indicated the formation of approximately spherical nanoparticles with slight agglomeration (Figure 3d).

#### 3.1.7. Attenuated Total Reflectance -Fourier Transform Infrared spectroscopy

ATR-FTIR (attenuated total reflectance – Fourier transform infrared spectroscopy) was conducted to identify functional groups in *L. camara* leaf extract as well as in the synthesized LCLE-Ag NPs. The major intense peaks in the leaf extract were observed approximately around 1000 cm^−1^, 1200 cm^−1^ 1700 cm^−1^, 2900 cm^−1^ and 3000 cm^−1^. The firm, intense broad bands at 2921.62 cm^−1^ and 2853.67 cm^−1^ suggest the presence of amine salts and alkane groups in LCLE-Ag NPs, respectively (Figure 4a). The small peak around 2500 cm^−1^ is the characteristic peak of the nitrile group and alkyne in the nanoparticles. Another intense band between 1500 cm^−1^-1750 cm^−1^ is attributed to a carbonyl group (41). The peak signifying carbonyl group was found to be weaker in synthesized LCLE-Ag NPs when compared with *L. camara* leaf extract (Figure 4b). This difference suggests participation of the carbonyl groups present in the *L. camara* leaf extract along with the silver during the nanoparticle formation. The phenol groups present in the leaf extract, around 1300 cm^−1^, are likely to be responsible for the bio-reduction of silver ions (48). The medium stretching of alkane and aldehyde, along with minor stretching of nitrile group and alcohols or ethers, indicates the presence of different phytoconstituents of *L. camara* in the nanoparticles, confirming the green synthesis.

**Fig 4.**
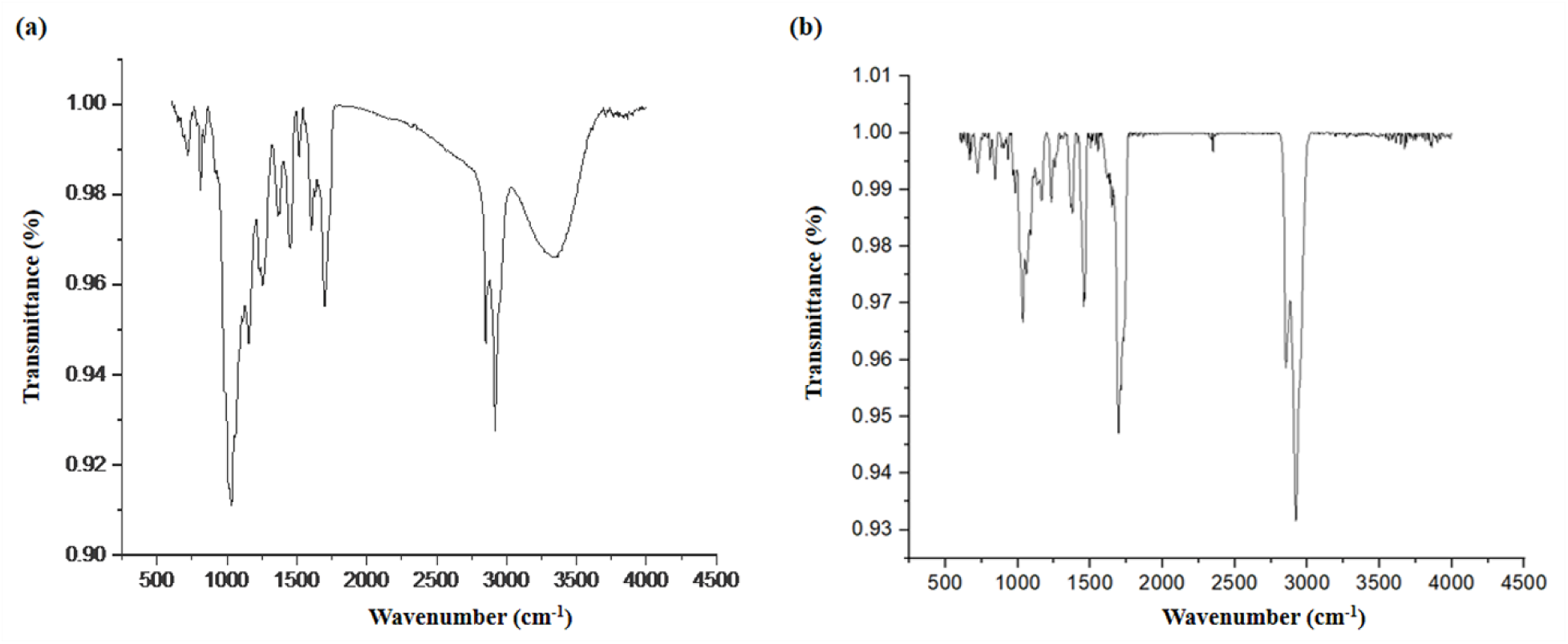
FTIR spectrum of (a) *Lantana camara* leaf extract, (b) Green synthesized LCLE-Ag NPs

### 3.2. Antibacterial effect of LCLE-Ag NPs against *Pseudomonas aeruginosa*

The minimum inhibitory concentration of prepared LCLE-Ag Np was checked at 5-10-20-50-100 μg/mL concentrations against *Pseudomonas aeruginosa* KPW.1-S1 and HRW.1-S3 (Figure 5a). For KPW.1-S1, at lower concentrations (5 μg/mL and 10 μg/mL), the nanoparticles exhibited non-significant effects, with only slight reductions in bacterial growth. Further, at 20 μg/mL, the nanoparticles achieved a significant reduction by 0.25-fold, whereas 50 μg/mL concentration of nanoparticles resulted in a significant decrease by 0.996-fold. In the presence of 100 μg/mL, bacterial growth was entirely inhibited, confirming the effectiveness of LCLE-Ag NPs at this dose.

**Fig 5.**
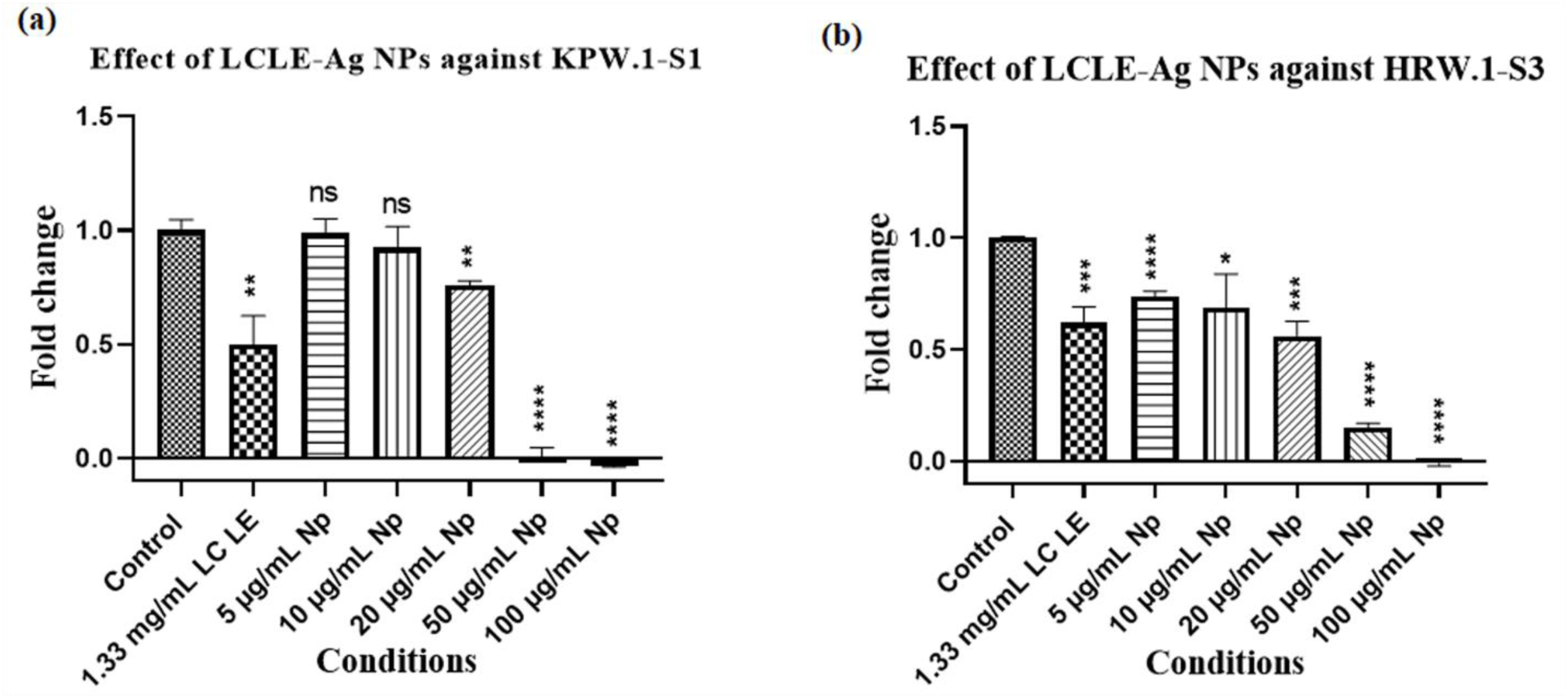
Minimum inhibitory concentration assay of green synthesized LCLE-Ag NPs against *P. aeruginosa* (a) KPW.1-S1 and (b) HRW.1-S3; Data presented as the mean ± SEM; n=9; N=3; *, P ≤ 0.05, **P ≤0.01, ***P ≤0.001, ****P ≤0.0001, while ‘ns’ indicates non-significance

Similarly, when the minimum inhibitory concentration assay was performed on *Pseudomonas aeruginosa* HRW.1-S3 (Figure 5b), it was found that at the lower doses of nanoparticles (5-10-20 μg/mL), the LCLE-Ag NPs showed a 0.27-fold, 0.32-fold, and 0.44-fold effect on bacterial growth, respectively, which further increased to 0.85-fold in the presence of 50 μg/mL LCLE-Ag NPs. At the highest dose of 100 μg/mL LCLE-Ag NPs, the bacterial growth was inhibited entirely, confirming the efficacy of LCLE-Ag NPs at this dose.

Therefore, the MIC concentration of the LCLE-Ag NPs for both the strains was determined to be 100 μg/mL.

For both the strains, 50-100 μg/mL concentrations of LCLE-Ag NPs showed more effect as compared to 1.33 mg/mL of LCLE, which is the maximal possible concentration of LCLE in the prepared nanoparticle (Fig. 5a,b).

### 3.3. Antibiofilm effect of LCLE-Ag NPs against *Pseudomonas aeruginosa*

The Biofilm load assay was performed to study the effectiveness of the prepared LCLE-Ag NPs against *Pseudomonas aeruginosa* KPW.1-S1 and HRW.1-S3 biofilm. Interestingly, for KPW.1-S1, it was observed that when maximal possible concentration of LCLE (1.33 mg/mL) was used, it encouraged biofilm formation significantly by a 4.85-fold compared to the untreated control (Figure 6a). Importantly, it was observed that the prepared nanoparticles could reduce the biofilm with increasing doses. In the presence of 10 μg/mL LCLE-Ag NP, a non-significant decrease was seen but at 20 μg/mL concentration of LCLE-Ag NPs, the biofilm load decreased by 0.75-fold compared to the control. Higher concentration (50 μg/mL) of LCLE-Ag NP further reduced the biofilm by 0.67-fold and at the concentration of 100 μg/mL, further reduction in biofilm formation was observed, with a 0.97-fold decrease relative to the control (Figure 6a).

**Fig 6.**
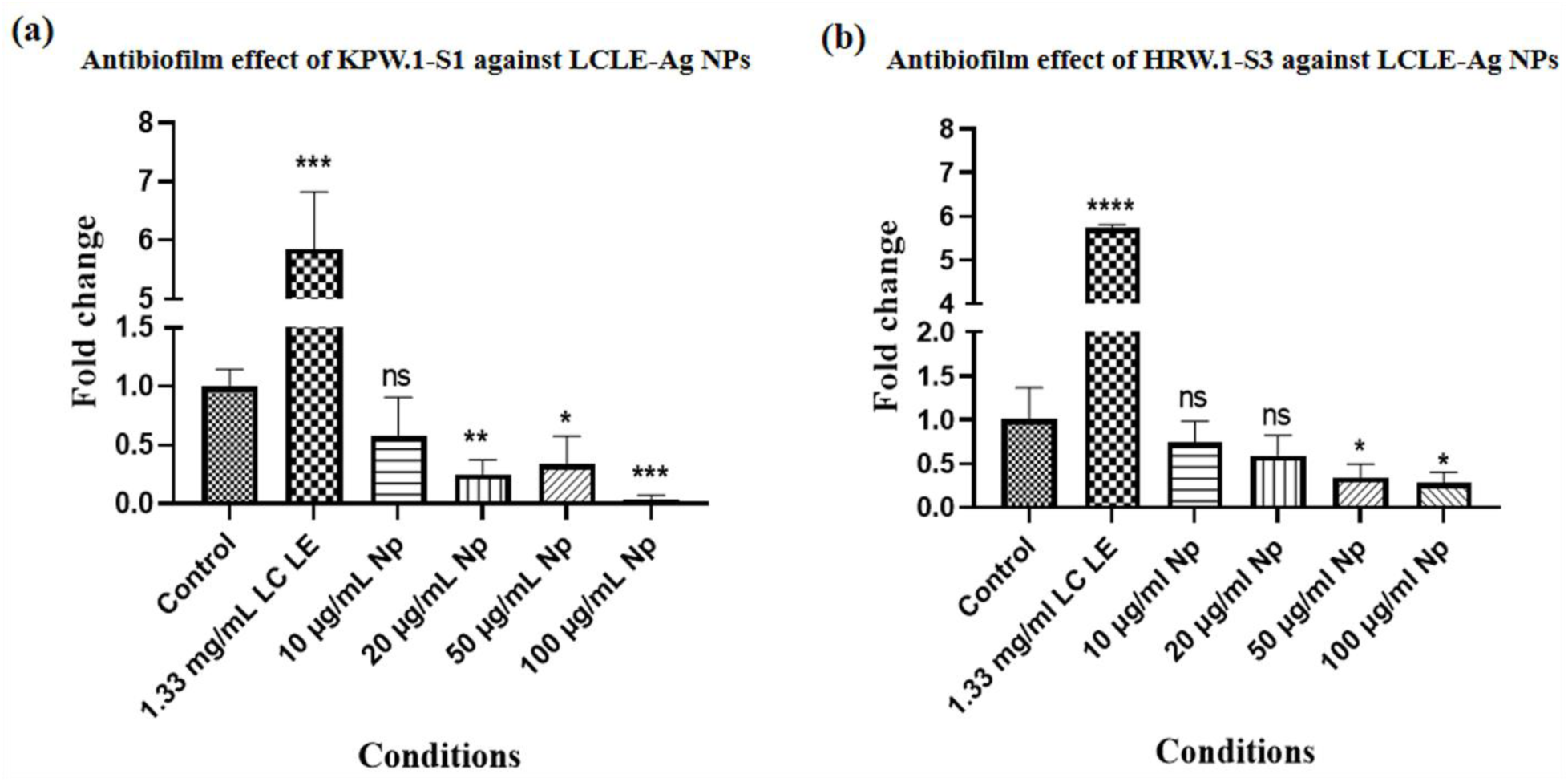
Biofilm load assay of *P. aeruginosa* KPW.1-S1 and HRW.1-S3 in the presence of LCLE-Ag NP (a) O.D. of biofilm cells of KPW.1-S1 (b) O.D. of biofilm cells of HRW.1-S3; Data represent mean ± SEM of n=9; N=3; *, P ≤ 0.05, **P ≤0.01, ***P ≤0.001, ****P ≤0.0001, while ‘ns’ indicates non-significance

The biofilm load assay was also performed against *Pseudomonas aeruginosa* HRW.1-S3 (Figure 6b). Relative to control, the maximal possible concentration of 1.33 mg/mL LCLE showed a significant 4.73-fold increase in biofilm load. At the lower doses of 10 μg/mL and 20 μg/mL, the nanoparticles had a non-significant effect on the biofilm, but at higher doses of 50 μg/mL and 100 μg/mL, the biofilm load was significantly decreased to 0.65-fold and 0.73-fold, respectively (Figure 6b). The images of the crystal violet-stained biofilms are given in Supplementary Figure S2 as a supporting document.

Thus, in both the strains the leaf extract at its maximal concentration can induce biofilm in comparison to control but the nanoparticle were effectively reducing biofilm at concentration of 50 μg/mL and 100 μg/mL.

### 3.4. Effect of LCLE-Ag NPs on the architecture of *Pseudomonas aeruginosa* biofilms

To visualize the reduction in the biofilm load and alterations in biofilm structures in the presence of LCLE-Ag NPs, confocal laser scanning microscopy (CLSM) and scanning electron microscopy (SEM) were performed.

#### 3.4.1. Confocal Laser Scanning Microscopy analysis

CLSM analysis showed the inhibitory effect of LCLE-Ag NPs on biofilm formation by PA strains (Figure 7). In comparison to the control samples, thick biofilm formation was observed in the presence of 1.33 mg/mL *L. camara* leaf extract. Interestingly, the biofilm formation was found to be decreased with the increasing concentrations of prepared LCLE-Ag NPs for both of the PA strains (Figure 7a, b).

**Fig 7.**
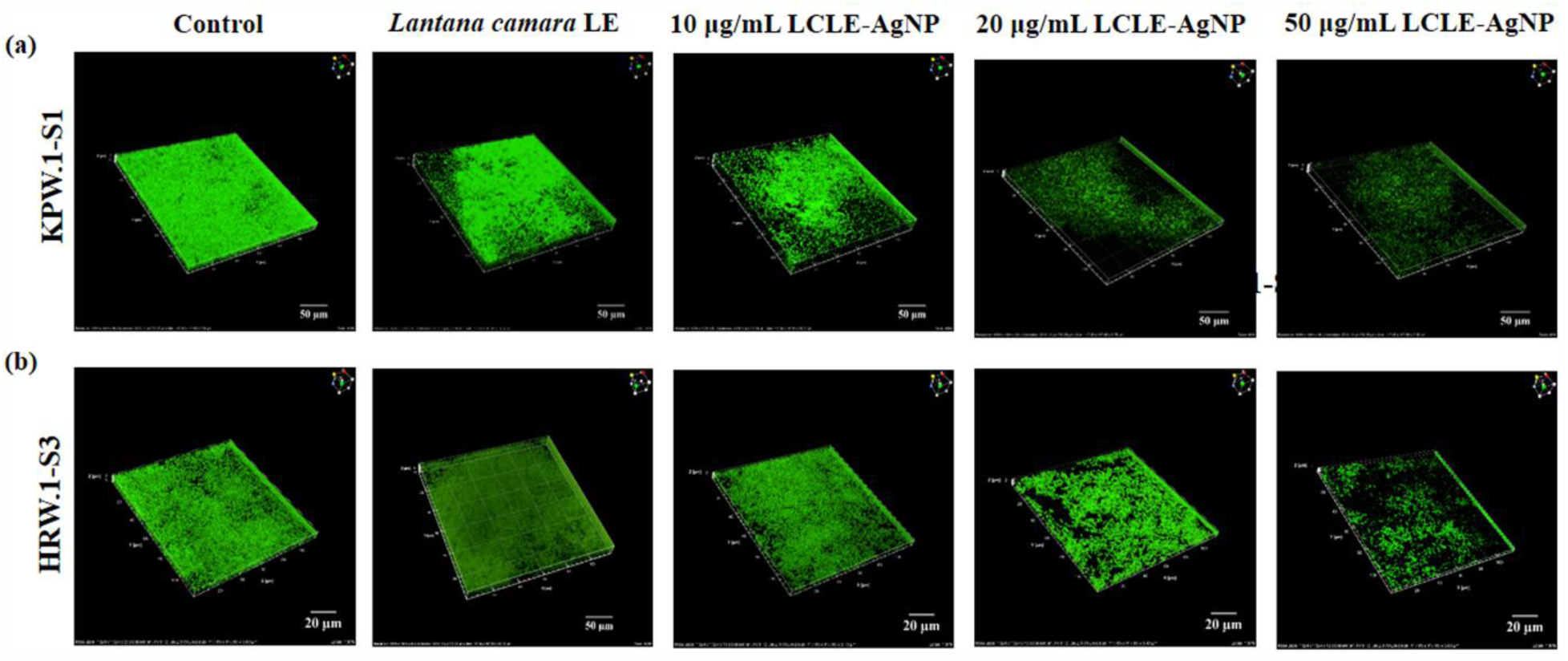
Representative confocal laser scanning microscopy images of *P. aeruginosa* biofilm in the presence of LCLE-Ag NP (a) 100X images of KPW.1-S1 biofilm (b) 100X images of HRW.1-S3 biofilm; Data represent mean ± SEM of n=9; N=3; *, P ≤ 0.05, **P ≤0.01, ***P ≤0.001, ****P ≤0.0001

#### 3.4.2. Scanning Electron Microscopy analysis

The SEM was performed to study the structure of the PA biofilms. For KPW.1-S1 (Figure 8a), it was observed that most of the cells in control samples adhered to the surface and with each other to form a dense biofilm. Treatment with 1.33 mg/mL *L. camara* leaf extract induced the bacterial cells to form a thick and robust biofilm with increased presence of extracellular polymeric substances and increased number of canopies. However, with the introduction of increasing concentrations of prepared nanoparticles, it was observed that the biofilm thickness decreased significantly. In the presence of 50 µg/mL of nanoparticles, mostly individually scattered cells were present with no substantial EPS production, whereas in the presence of 100 µg/mL of nanoparticles, very few individual cells, along with some cell debris, were visible (Figure 8a).

**Fig 8.**
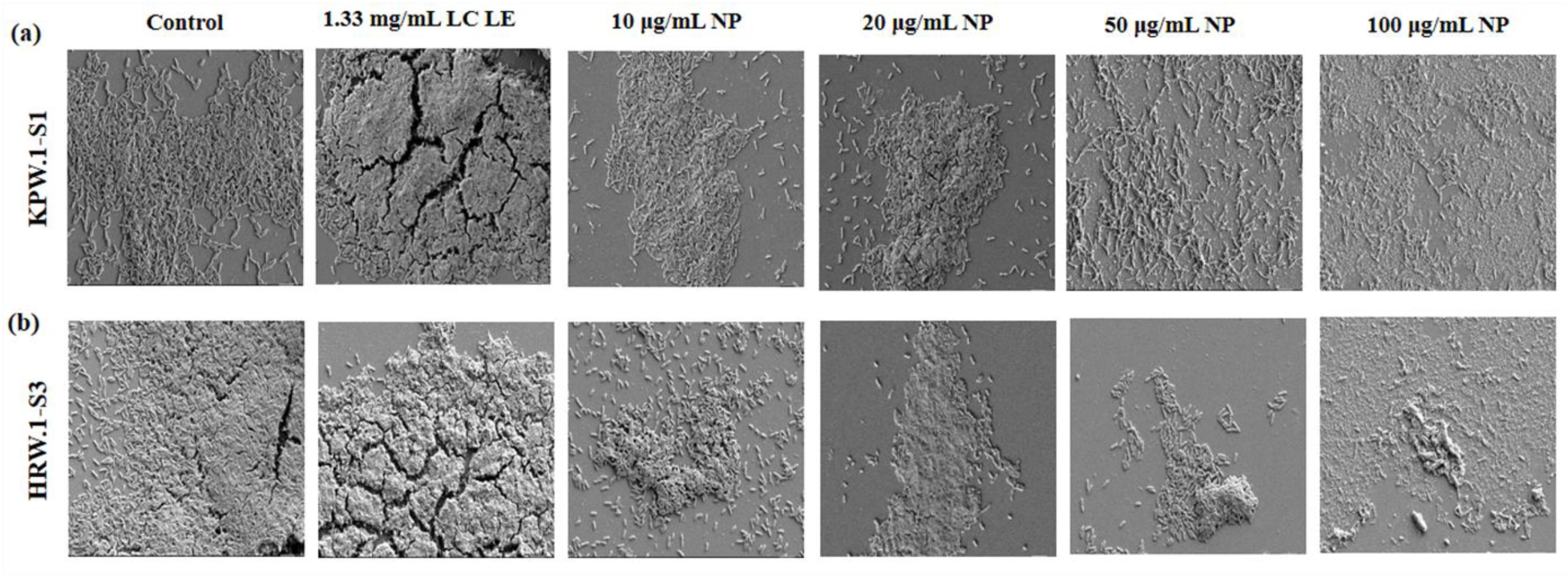
Representative Scanning Electron Microscope images of *P. aeruginosa* biofilm in the presence of LCLE-Ag NP (a) 5.0KX images of KPW.1-S1 biofilm (b) 5.0KX images of HRW.1-S3 biofilm; Data represent mean ± SEM of n=6; N=2; *, P ≤ 0.05, **P ≤0.01, ***P ≤0.001, ****P ≤0.0001

In the case of HRW.1-S3 (Figure 8b), the control condition showed the formation of a thick biofilm. The presence of LCLE alone further increased the biofilm formation, with a more thickened biofilm structure with an increased number of canopies. The introduction of low doses of nanoparticles decreased the overall biofilm formation in comparison to the control and LCLE alone. The presence of 50 µg/mL showed a few clumped cells, and in the highest dose (100 µg/mL), very few cells, along with cell debris, were visible (Figure 8b).

### 3.5. Visualization of Extracellular polysaccharide and extracellular DNA analysis

The alterations in the abundance of extracellular polysaccharides and extracellular DNA (eDNA) due to nanoparticles treatment were examined using dual staining of biofilms with Propidium iodide (PI) and Concanavalin A (Con A). PI binds to extracellular DNA contributed by secretion or membrane-compromised cells present in the matrix, resulting in characteristic red fluorescence, and Con A binds to mannose residues of the extracellular polysaccharides in the biofilm matrix, indicated by blue fluorescence, thus differentiating both the major components of the biofilm matrix. In case of PA KPW.1-S1 biofilm, Con A intensity decreased by 0.10-fold and 0.24-fold in comparison to control at lower doses (10 µg/mL and 20 µg/mL) of LCLE-Ag NPs respectively. At 50 µg/mL dose of the nanoparticles, the Con A intensity decreased by 0.52-fold, suggesting very less secretion of polysaccharides at the higher dose of LCLE-Ag NPs. Correspondingly, at 10 µg/mL and 20 µg/mL, PI intensity reduced by respective 0.27-fold and 0.68-fold, which further decreased significantly at higher doses (50 µg/mL) by 0.88-fold (Figure 9c).

**Fig 9.**
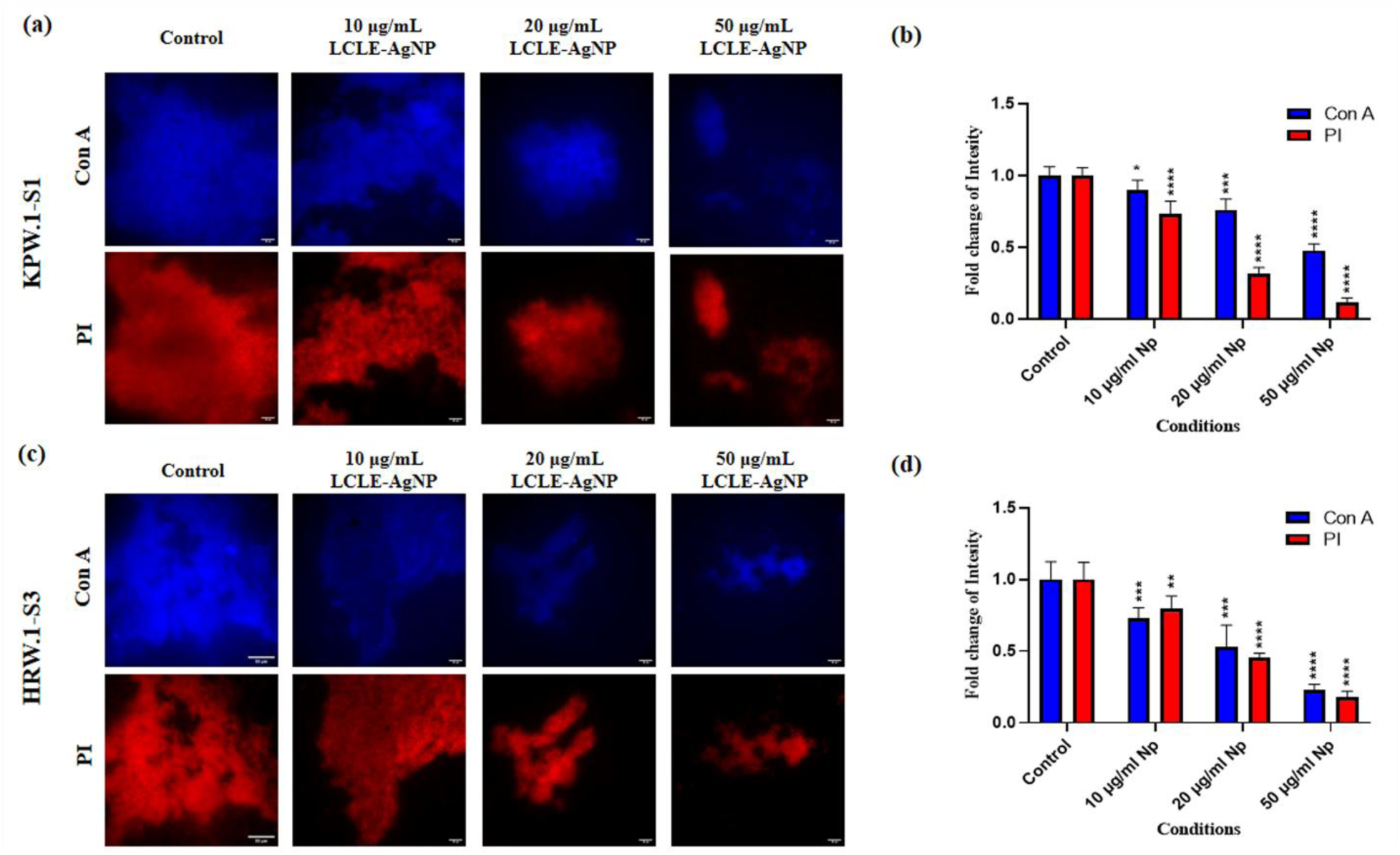
Representative epifluorescence images of *P. aeruginosa* biofilm stained with Con A (blue) and PI (red) in the presence of LCLE-Ag NP (a) 60X images of KPW.1-S1 biofilm (b) fold change intensity for the strain KPW.1-S1 (c) 60X images HRW.1-S3 biofilm (d) fold change intensity for the strain HRW.1-S3; Data represent mean ± SEM of n=6; N=2; *, P ≤ 0.05, **P ≤0.01, ***P ≤0.001, ****P ≤0.0001

Analyses of both the exopolysaccharide and eDNA components of the PA KPW.1-S1 biofilm matrix showed a significant decrease with the treatment of increasing concentrations of nanoparticle (Figure 9a). In the case of HRW.1-S3 strain, a similar trend was found in eDNA and extracellular polysaccharides (Figure 9c). The reduction in the extracellular components supports the SEM observations, where substantial decrease of EPS was visible. This suggests that the nanoparticle significantly reduces the secretion of biofilm matrix components resulting in an overall decrease in biofilm formation.

Similarly, for the PA HRW.1-S3, Con A fluorescence intensity decreased significantly, showing a 0.27-fold reduction at 10 µg/mL, 0.47-fold reduction at 20 µg/mL, and a 0.77-fold reduction at 50 µg/mL of nanoparticles. PI intensity also declined by 0.23-fold at 10 µg/mL, with further reductions of 0.54-fold and 0.82-fold observed at 20 µg/mL and 50 µg/mL, respectively (Figure 9d). These observations indicate that LCLE-Ag nanoparticles effectively weaken the biofilm matrix by diminishing the secretion of major EPS components.

### 3.6. Effect of LCLE-Ag NPs on the preformed biofilm of *Pseudomonas aeruginosa*

Biofilm eradication assays were performed against *Pseudomonas aeruginosa* KPW.1-S1 and HRW.1-S3 to study the effectiveness of LCLE-Ag NPs on the preformed biofilm. For preformed biofilm in KPW.1-S1, addition of LCLE further increased the biofilm by 5.41-fold, whereas the LCLE-Ag NPs showed a decrease in the formed biofilm. Although there was no significant decrease in the presence of lower doses (10 µg/mL and 20 µg/mL) of nanoparticle, higher doses of nanoparticles, (50 µg/mL and 100 µg/mL) could reduced the formed biofilm by 0.71-fold and 0.82-fold, respectively, showcasing its ability to eradicate biofilm (Figure 9a).

In the case of HRW.1-S3, the 1.33 mg/ml LCLE showed a 5.62-fold increase in biofilm load. The lower doses of nanoparticles (10 µg/mL and 20 µg/mL) resulted in a non-significant decrease in the formed biofilm, whereas the higher doses of nanoparticles (50 µg/mL and 100 µg/mL) reduced the formed biofilm significantly by 0.61-fold and 0.67-fold (Figure 9b).

The images of the crystal violet-stained biofilm are given in Supplementary Figure S3 as a supporting document.

### 3.7. Morphological study on the effect of LCLE-Ag NPs on Gentamicin induced preformed biofilm

The confocal microscopy was performed to study the efficacy of the nanoparticles against preformed biofilms induced by gentamicin in PA strains KPW.1-S1 and HRW.1-S3 (Figure 11 a,b). In PA KPW.1-1, the lower doses of 10 μg/mL and 20 μg/mL LCLE-Ag NPs showed reduction in the biofilm thickness compared to the control biofilm. The higher doses of nanoparticles, 50,100-μg/mL, reduced the biofilm mass as well as the thickness decreased significantly (Figure 11a).

**Fig 10.**
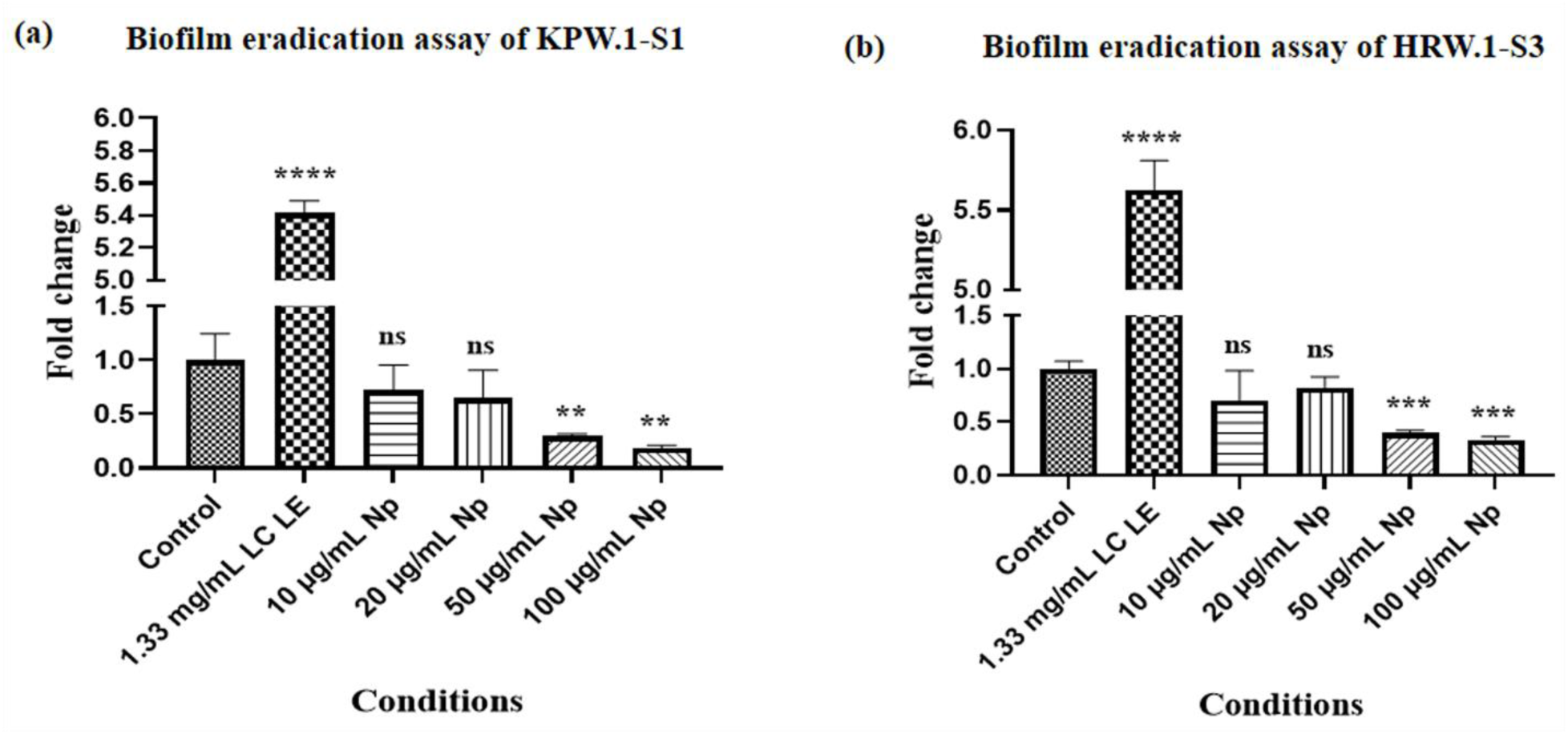
Biofilm eradication assay of *P. aeruginosa* KPW.1-S1 and HRW.1-S3 in the presence of LCLE-Ag NP (a) O.D. of biofilm cells of KPW.1-S1 (b) O.D. of biofilm cells of HRW.1-S3; Data represent mean ± SEM of n=9; N=3; *, P ≤ 0.05, **P ≤0.01, ***P ≤0.001, ****P ≤0.0001, while ‘ns’ indicates non-significance

**Fig 11.**
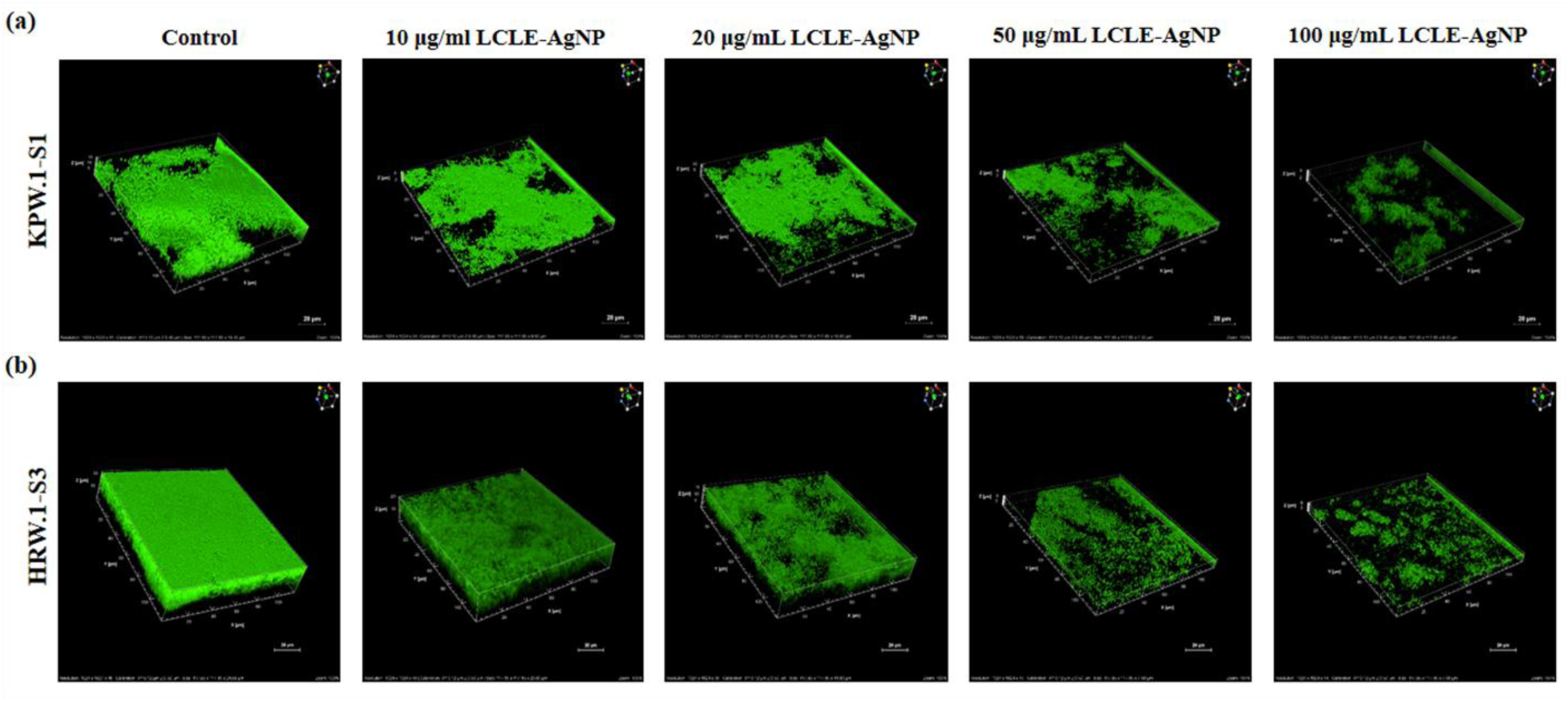
Representative confocal laser scanning microscopy images of gentamicin induced *P. aeruginosa* KPW.1-S1 and HRW.1-S3 biofilm in the presence of LCLE-Ag NP (a) 100X images of KPW.1-S1 biofilm (b) 100X images of HRW.1-S3 biofilm; Data represent mean ± SEM of n=6; N=2; *, P ≤ 0.05, **P ≤0.01, ***P ≤0.001, ****P ≤0.0001

The PA strain HRW.1-S3 also showed significant reduction in biofilm in the presence of higher doses of the nanoparticle indicating that the nanoparticle can effectively reduce biofilm formed due to the sub-MIC dose of gentamicin (Figure 11b).

### 3.8. Quantitative analysis of ROS production in P. aeruginosa biofilms in the presence of nanoparticle

The ROS production was estimated in *Pseudomonas aeruginosa* KPW.1-S1 and HRW.1-S3 biofilms in the presence and absence of LCLE-Ag NPs using fluorescence microscopy. It was observed that the lower doses (10 and 20-μg/mL) of LCLE-Ag NPs could increase the ROS production by 5.21-fold and 6.88-fold, respectively, as compared to control condition. However, at a higher dose (50 μg/mL) of LCLE-Ag NPs, the ROS production increased only by 3.85-fold, possibly due to high cell death in the given condition (Figure 12a,c).

**Fig 12.**
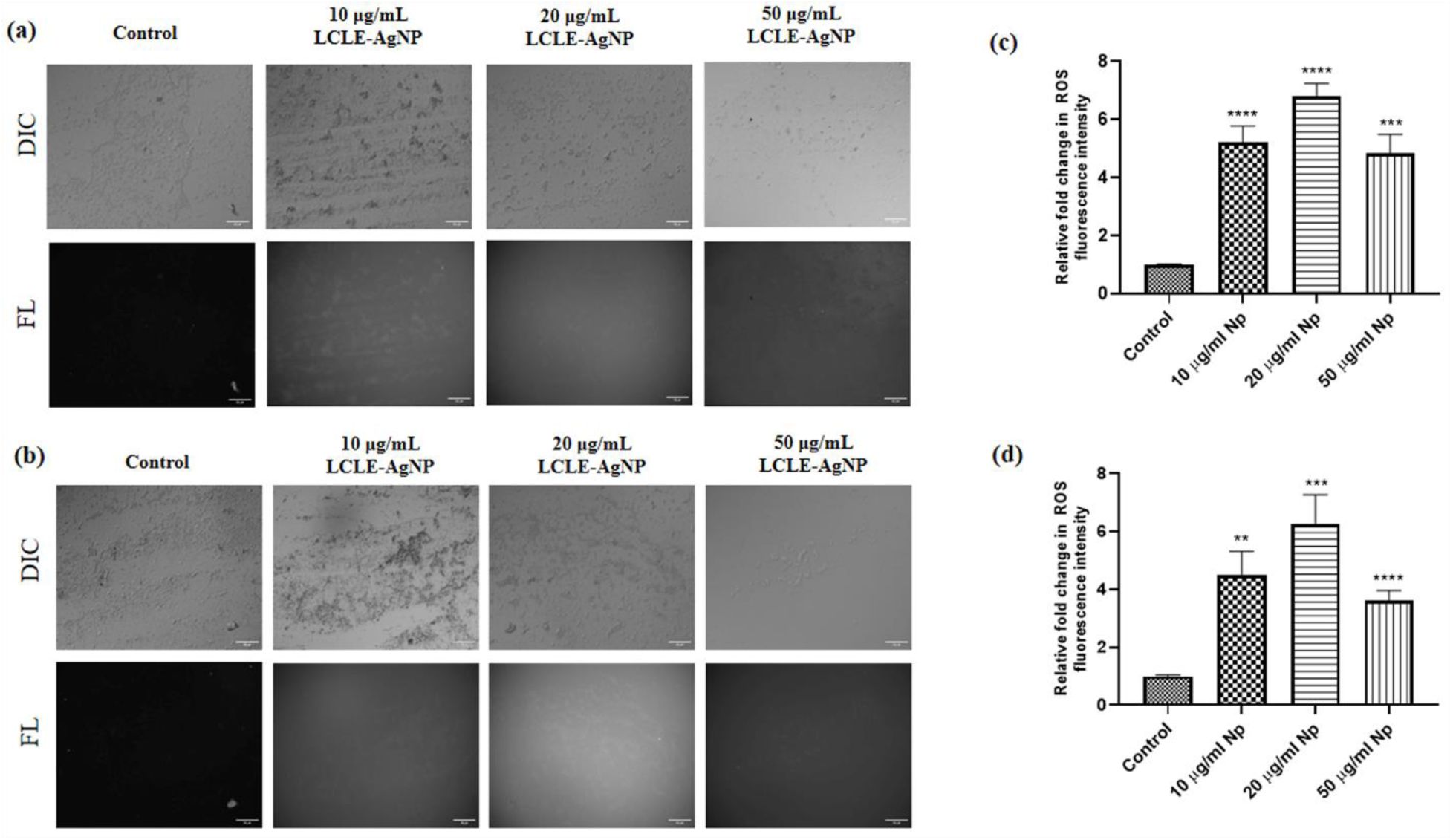
Representative epifluorescence images ROS generation in *P. aeruginosa* KPW.1-S1 and HRW.1-S3 biofilm cells in the presence of LCLE-Ag NP (a) 40X images of KPW.1-S1 biofilm (b) 40X images of HRW.1-S3 biofilm (c) relative fold change for the strain KPW.1-S1 (d) relative fold change for the strain HRW.1-S3; Data represent mean ± SEM of n=6; N=2; *, P ≤ 0.05, **P ≤0.01, ***P ≤0.001, ****P ≤0.0001

In the case of the HRW.1-S3 strain, at the lower doses of LCLE-Ag NPs (10 μg/mL and 20 μg/mL), the ROS production was increased by 4.49 and 6.24 folds, respectively. Subsequently, at the higher dose of 50 μg/mL LCLE-Ag NPs, the ROS production increased only till 2.62-fold (Figure 12b,d) may be due to the same reason as mentioned in case of KPW.1-S1 strain.

### 3.9. Effect of LCLE-Ag NPs on *lasI* and *pqsA* expression in PA KPW.1-S1

As LCLE-Ag NPs showed a significant effect on formation and eradication of *Pseudomonas aeruginosa* biofilms, the expression of two important quorum sensing genes namely, *lasI* and *pqsA* were analyzed.

Both the *lasI* and *pqsA* expressions were found to be decreasing with increasing doses of LCLE-Ag NPs (Figure 13a). In case of *lasI*, the lower dose (10 μg/mL) of nanoparticles showed a non-significant decrease, while the higher doses of 20 μg/mL and 50 μg/mL LCLE-Ag NPs showed 0.52-fold and 0.81-fold decreases, respectively (Figure 13b). Similarly, *pqsA* expression showed a significant decrease of 0.37-fold, 0.61-fold, and 0.86-fold in the presence of 10 μg/mL, 20 μg/mL, and 50 μg/mL LCLE-Ag NPs, respectively (Figure 13c).

**Fig 13.**
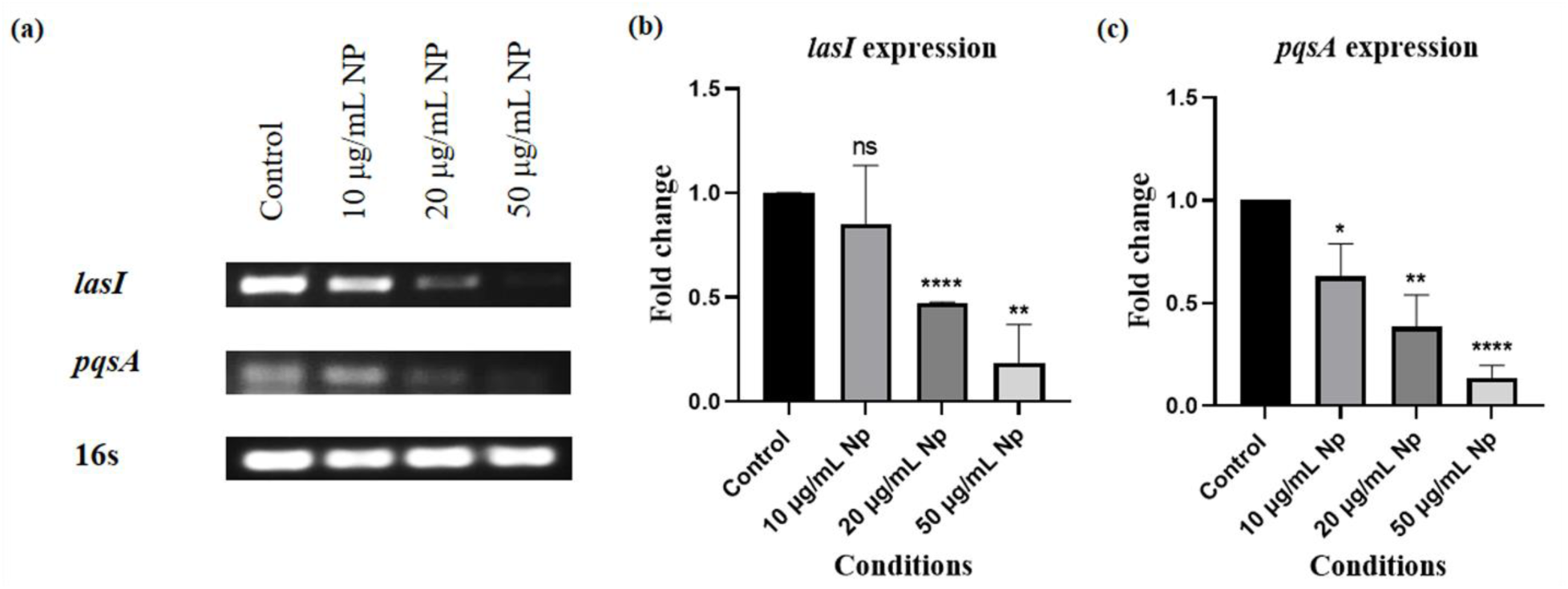
Representative images of semi-quantitative PCR amplified DNA of respective genes in agarose gel (a) agarose gel images of *lasI, pqsA* and 16s (b) fold change in *lasI* expression (c) fold change in *pqsA* expression; Data represent mean ± SEM of n=6; N=2; *, P ≤ 0.05, **P ≤0.01, ***P ≤0.001, ****P ≤0.0001

Overall, gene expression analysis of *lasI* and *pqsA* showed a decrease in expression pattern with increasing LCLE-Ag NP doses, indicating disruption of quorum-sensing pathways which are essential for biofilm formation.

### 3.10. Effect of LCLE-Ag NPs on the viability of HEK293T cells

The cytotoxic effect of different doses of LCLE-Ag NPs and their equivalent doses of silver ions was studied in HEK293T cells. For all the doses, LCLE-Ag NPs could not induce any significant cytotoxicity in HEK293T cells as compared to the untreated cells (Figure 10). Whereas the equivalent doses of silver showed significant cytotoxicity in the HEK293T cells. These results clearly indicate that normal silver ions are toxic to cells, but the green synthesized LCLE-Ag NP has no significant toxicity on the HEK293T cells, making it as a potent candidate for medicinal uses.

## 4. Discussion

In the present study, silver nanoparticles were successfully green-synthesized using *Lantana camara* leaf extract, which has emerged as an eco-friendly and sustainable approach worldwide. The phytochemical constituents of the plant extracts reduced silver ions (Ag^+^ to Ag^0^) and stabilized the nanoparticles by acting as a capping agent. Previous studies have shown that bioactive compounds present in the plant extract acts both as a reducing and a capping agent (49,50).

UV-visible spectroscopy serves as a key analytical tool for determining the formation and assessing the stability of metal nanoparticles in aqueous solution with a distinct spectrum. In case of our study, the successful formation of LCLE-Ag NPs was characterized by a surface plasmon resonance (SPR) peak observed between 350-470 nm (Figure 1). Such SPR signatures are consistent with previously reported phytogenic silver nanoparticles (51). Zeta potential and DLS results indicated the good stability of the nanoparticle, with values indicating strong electrostatic repulsion (-9.54 mV) and a particle size range of 30-50 nm (Figure 2). This suggests that the phytochemicals present in the leaf extract are enhancing the repulsion between negatively charged particles and preventing agglomeration as stated in previous reports (52,53).

While evaluating the morphology of LCLE-Ag NPs, the SEM analysis revealed the size of nanoparticles to be 25-35 nm, having a spherical morphology with slight clustering, which was further confirmed by TEM and AFM analysis (Figure 3). This clustering is normal due to some factors, such as concentration and environmental conditions, despite their stabilizing effects (44,45). EDX analysis confirmed the elemental composition of the synthesized LCLE-Ag NPs. The EDX spectrum for *L. camara* leaf extract showed a weight percentage of 0% for Ag, indicating that the amount of silver in nanoparticles is not from *L. camara* but due to the successful formation of nanoparticles. The ATIR-FTIR study on chemical composition further confirmed the presence of several functional groups and suggests their involvement in the formation of nanoparticles (Figure 4). The spectra showed a strong signal for phytoconstituents, employing the successful formation of green synthesized silver nanoparticles. The FTIR spectra revealed prominent peaks corresponding to hydroxyl (-OH), carbonyl (C=O), and amine (N-H) groups, which are indicative of the phytochemical constituents from *L. camara* acting as reducing and stabilizing agents. These functional groups likely facilitate the capping of nanoparticles, enhancing their stability and bioactivity as similarly shown in other green nanoparticle studies (46,47).

The LCLE-Ag NPs exhibited strong antibacterial, antibiofilm, and biofilm-eradicating activity against PA strains KPW.1-S1 and HRW.1-S3 in a dose-dependent manner.

For the KPW.1-S1 strain, the lower concentrations of the nanoparticles did not result in any significant reduction in growth. In contrast, the higher nanoparticle doses (50 µg/mL and 100 µg/mL) caused a marked and significant inhibition of bacterial growth. For the HRW.1-S3 strain, all tested concentrations of LCLE-Ag NPs exhibited significant antibacterial activity, with the higher doses exerting a more antibacterial effect than LCLE alone (Figure 5). Such differences in inhibitory potentials between the nanoparticle and leaf extract at elevated NP doses strongly support the superior efficiency of the nanoparticles over the crude extract and the probable reason for this antibacterial activity can be attributed to the disruption of the cell membranes, ROS generation, and impact on overall cell functions as seen in various other studies (57–59).

The biofilm load assay for both the strains further demonstrated that the synthesized nanoparticle had an inhibitory effect on the biofilm load in a dose-dependent manner, whereas the maximal possible concentration of LCLE rather promoted biofilm formation (Figure 6). Earlier reports confirmed the biofilm-disruptive properties of AgNPs synthesized using *A. cobbe* leaf extract but did not explore the biofilm-inducing effects of the corresponding plant extracts (60). Our findings, therefore, significantly extend the existing knowledge by uncovering that *L. camara* leaf extract alone may induce PA biofilm formation, thus highlighting the importance of LCLE-Ag NPs for both prevention and eradication of resilient bacterial communities.

This effect of LCLE-Ag NPs on the biofilm was further confirmed when we visualized the biofilm using CLSM and SEM. The CLSM imaging supported the observation that LCLE alone induced 3D biofilm formation, but LCLE-Ag NPs could inhibit the biofilm (Figure 7). Consistent with this, SEM analysis showed a substantial reduction in biofilm biomass and visibly diminished EPS in nanoparticle-treated samples of both PA strains, while the leaf extract alone enhanced biofilm structure, producing distinct canopy-like formations (Figure 8). Con A-PI staining confirmed the significant reduction of two extracellular polymeric components: exopolysaccharides and extracellular DNA, which are important for biofilm formation (44,61) in the presence of the nanoparticles (Figure 9). These results in the decrease in EPS components correlated with the observed SEM results (Figure 8). Siddique and his coworkers primarily demonstrated that silver nanoparticles suppress EPS production (62); however, the study did not investigate EPS subcomponents or 3D biofilm morphology in depth. Here, we not only confirmed biomass loss but also quantified matrix-specific disruption, offering mechanistic evidence for nanoparticle-induced biofilm reduction. Further, the biofilm eradication potential of the nanoparticles were assessed on preformed biofilm as well as on gentamicin induced preformed biofilm. Gentamicin is a commonly prescribed drug against PA, and previous findings showed that it can induce biofilm formation by PA in the presence of sub-MIC doses (42,63). The LCLE-Ag NPs were found to be able to reduce preformed biofilms, whereas leaf extract alone further enhanced the biofilm, as evident by the biofilm load assay (Figure 10). The effect of nanoparticles on gentamicin-induced biofilm was further studied using CLSM to understand how effectively nanoparticles can reduce it. The results of the CLSM studies showed that the higher doses of nanoparticles were effectively able to minimize the biofilm thickness and biomass compared to the control, suggesting the strong potential of nanoparticles in the therapeutic world (Figure 11). Tabassum et al. demonstrated the antibiofilm and biofilm eradication potential of Pyoverdine-silver nanoparticles (64). In continuation of these efforts, our study further evaluates the efficacy of nanoparticle against antibiotic-induced biofilms by demonstrating that LCLE-Ag NPs strongly disperse both conventional and gentamicin-induced biofilms, thereby reinforcing their therapeutic superiority. Previous studies have shown that ROS affects biofilms by degrading the biofilm matrix (65,66), and thereby inhibits biofilm formation (67). In the present study, the ROS intensity was found to be increased significantly in the presence of nanoparticles (Figure 12), however, the increase was slightly less in the higher dose of 50 µg/ml, probably due to early cell death as reported earlier (68). Hamida et al. demonstrated that silver nanoparticles synthesised using *Desertifilum sp.* exert their antimicrobial activity mainly through oxidative stress induced by ROS generation, leading to cell death (69). Notably, the effective concentrations of LCLE-AgNPs used in the present study was comparatively lower, as compared to the nanoparticles reported by Hamida et al., showcasing the better efficacy of the silver nanoparticle synthesized using LCLE.

A dose-dependent decrease in the expression of two major quorum-sensing genes*, lasI and pqsA,* in PA was observed upon treatment with LCLE-Ag NPs (Figure 13). Down-regulation of these genes indicates that the nanoparticles likely interfere with autoinducer synthesis in both the Las and PQS systems, resulting in decreased biofilm formation. Gurunathan et al. highlighted the phenotypic aspects of the antibacterial and antibiofilm effects of *A. cobbe* leaf extract mediated silver nanoparticles (60). Our findings further complement these findings by demonstrating that LCLE-Ag NPs modulate QS at the molecular level by directly downregulating QS-regulatory genes, thereby inhibiting biofilm formation.

Having addressed the bactericidal and antibiofilm properties of LCLE-Ag NPs, it was essential to assess the toxicity of LCLE-Ag NP on the mammalian cell line. The MTT assay of Human Kidney Epithelial-like cells with different doses of LCLE-Ag NPs exhibited minimal toxicity, with cell viability ranging from 90% to 100% (Figure 14). In contrast, equivalent doses of silver (with respect to LCLE-Ag NP) showed high toxicity. Thus, the comparative toxicity profiling reinforces that the green-synthesized nanoparticles offer a safer and favorable alternative over conventional silver ion.

**Fig 14.**
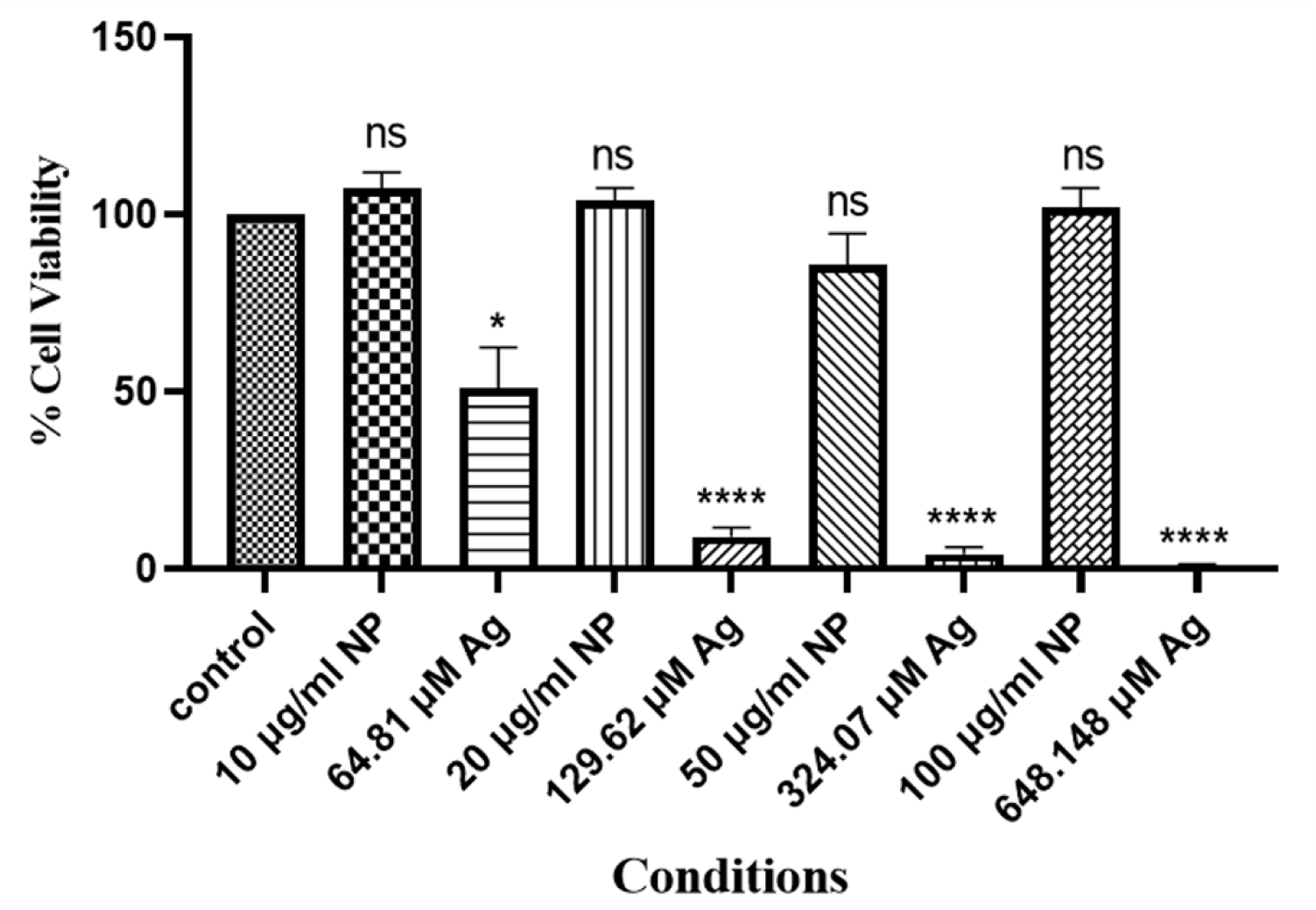
Quantification of viable cells by MTT assay in HEK293T cells in the presence of LCLE-Ag NPs and its equivalent silver concentration; Data represent mean ± SEM of n=9; N=3; *, P ≤ 0.05, **P ≤0.01, ***P ≤0.001, ****P ≤0.0001

Several studies have demonstrated the successful green-synthesis of silver nanoparticles and their efficacy against PA (70,71). Ajitha et al. reported the antibacterial potential of *L. camara* leaf, which exhibited positive effects against both Gram-negative and Gram-positive bacteria (72). In another study, Momani et al. demonstrated the antibiofilm activity of biosynthesized silver nanoparticles using *P. harmala* seed extract against PA isolates obtained from cystic fibrosis patients, highlighting the potential of plant-mediated nanoparticles (22), thus encouraged exploration of *L. camara* leaf extract as a biocompatible agent in this study. Our present work further advances this line of research by showing that L. camara-mediated silver nanoparticles are capable of exerting antibacterial and antibiofilm effects, including the inhibition of biofilm formation, as well as the eradication of conventional and antibiotic-induced biofilms against *Pseudomonas aeruginosa.* At the cellular level, this study demonstrated nanoparticle effects leading to cell damage/death by ROS production and molecular emphasis on quorum-sensing genes. Additionally, *L. camara*-mediated silver nanoparticles could not exert any significant cytotoxicity against Human Kidney Epithelial-like cells.

In conclusion, the enhanced efficacy and reduced toxicity of LCLE-Ag NPs synthesized using *L. camara* may be attributed to their nanoscale size, high surface area-to-volume ratio, ROS generation and their inhibitory effect on the quorum sensing systems. Future studies on elucidating the detailed molecular mechanism underlying the antibiofilm activity of the LCLE-Ag NPs will help to find relevant targets in *Pseudomonas* biofilms for remedial purposes.

## Acknowledgment

All authors would like to acknowledge the imaging facilities of the Indian Institute of Science Education and Research, Kolkata. The authors would also acknowledge Prof. Rahul Banerjee, DCS IISER Kolkata, for helping with the ATR-FTIR analysis. Authors also thank Arundhaty Pal and Sourav Sanyal, DBS IISER Kolkata for helping in MTT assays. SKS acknowledges IISER Kolkata for providing fellowship.

## CRediT authorship contribution statement

Conceptualization, **ADR, SKS and TKS;** Methodology**, ADR and SKS;** Investigation**, ADR and SKS;** Formal analysis**, ADR, SKS and TKS;** Data curation**, ADR and SKS;** Resources, **TKS;** writing—original draft preparation**, ADR and SKS;** writing—review and editing**, ADR, SKS and TKS;** supervision**, TKS;** funding acquisition**, TKS; All authors have read and agreed to the published version of the manuscript.**

## Data availability

All data generated or analyzed during this study are included in this present article.

## Ethical Approval

Not applicable.

## Consent to Participate

All authors had consented to participate in the study.

## Consent for Publication

All authors have given consent for publication.

