## Supplementary File for "Antimicrobial and Biofilm-Inhibitory Potential of Green Synthesized Silver Nanoparticles from *Lantana camara* against *Pseudomonas aeruginosa*"

### S1. Energy Dispersive X-ray spectroscopy analysis

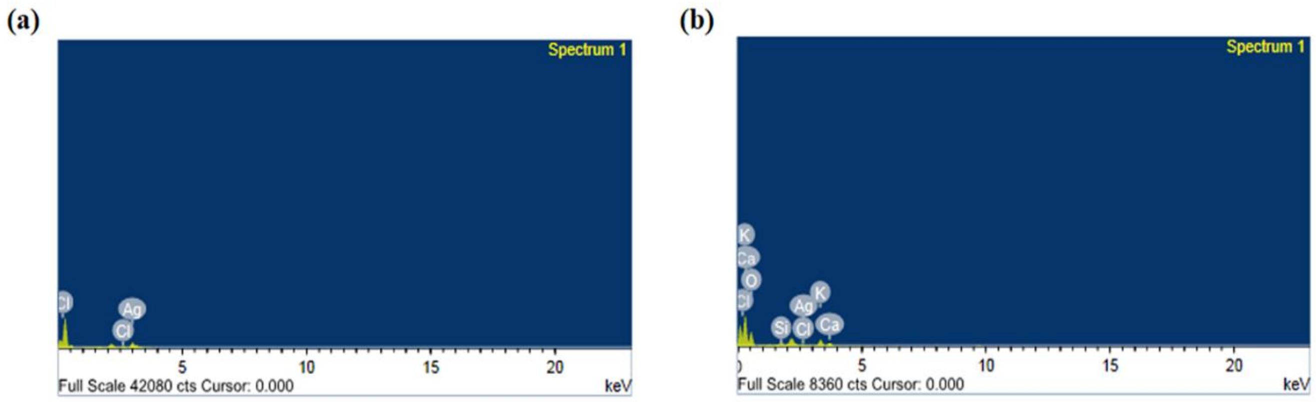


**Fig. S1.** EDX spectrum of (a) LCLE-Ag nanoparticles (b) *L. camara* leaf extract

**Figure S2. Estimation of biofilm load by crystal violet assay**

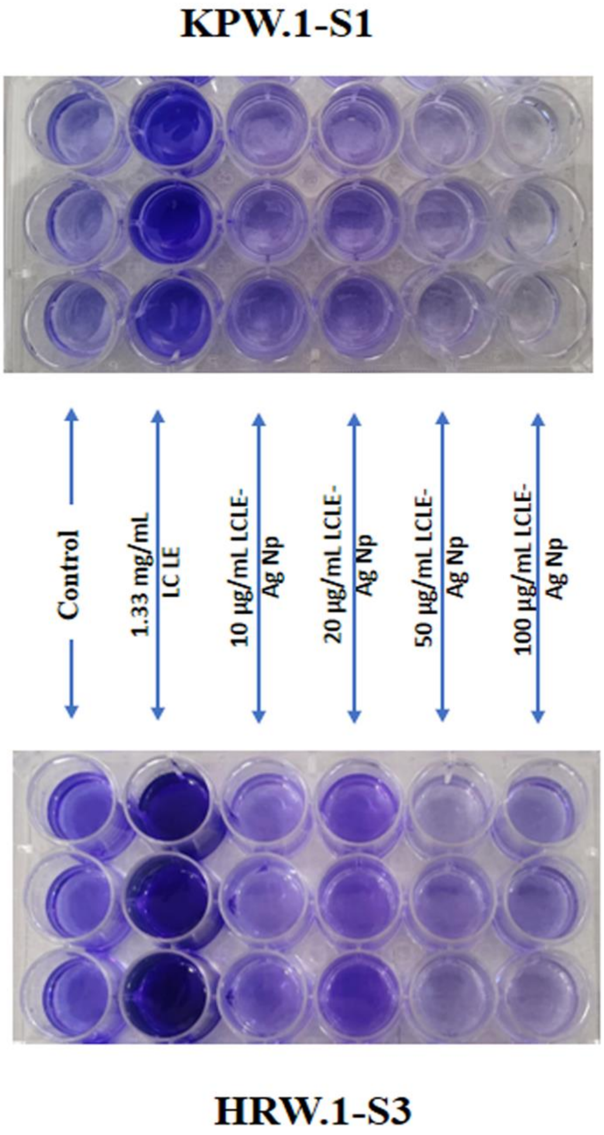


Fig. S2. Crystal violet staining of 24-well plates of KPW.1-S1 and HRW.1-S3 in different conditions

**Figure S3. Eradication of biofilm load by Nanoparticles**

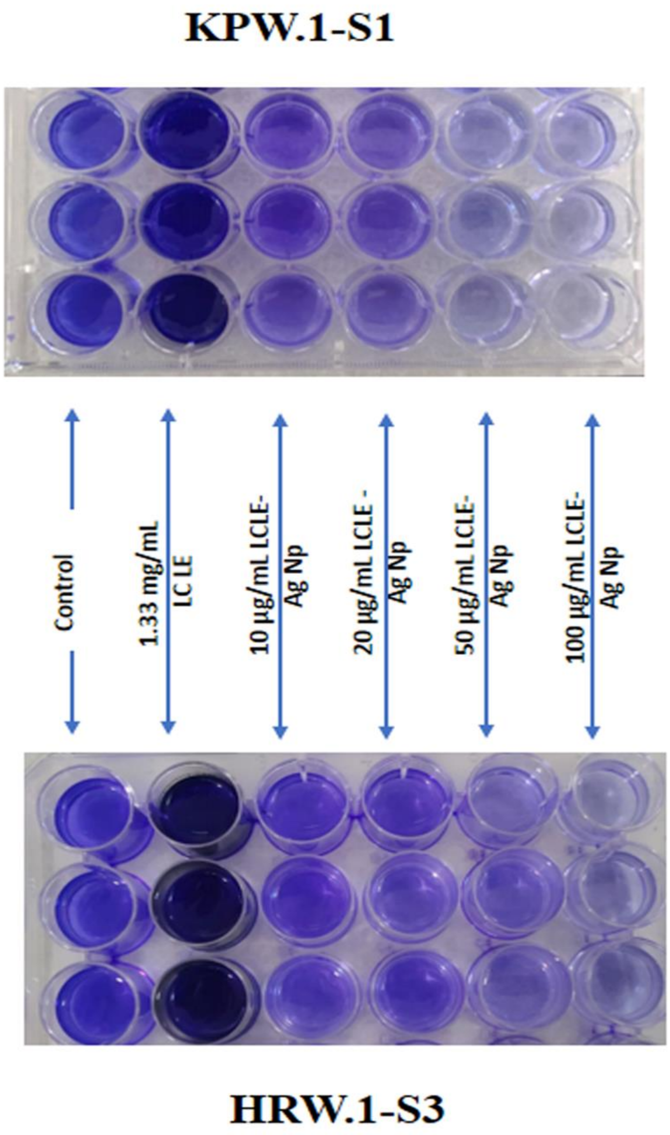


Fig. S3. Crystal violet staining of 24-well plates of biofilm eradication assay of KPW.1-S1 and HRW.1-S3 in different conditions
